# Rapid establishment of a tumor-retained state curtails the contribution of conventional NK cells to anti-tumor immunity in solid cancers

**DOI:** 10.1101/2023.08.10.552797

**Authors:** Isaac Dean, Colin Y.C. Lee, Zewen K. Tuong, Zhi Li, Christopher A. Tibbitt, Claire Willis, Fabrina Gaspal, Bethany C. Kennedy, Veronika Matei-Rascu, Rémi Fiancette, Caroline Nordenvall, Ulrik Lindforss, Syed Murtuza Baker, Christian Stockmann, Veronika Sexl, Gianluca Carlesso, Scott A. Hammond, Simon J. Dovedi, Jenny Mjösberg, Matthew R. Hepworth, Menna R. Clatworthy, David R. Withers

**Author notes:** Correspondence: Prof. David R Withers, Prof. Menna R. Clatworthy. **Abbreviations:** CAR, chimeric antigen receptor; cLN, contralateral LN; cNK, conventional Natural Killer; dLN, draining LN; ICB, immune checkpoint blockade; scRNA-seq, single cell RNA sequencing; TIL, tumor infiltrating lymphocyte; TME, tumour microenvironment; UMAP, Uniform Manifold Approximation and Projection.

## Abstract

Immune cell dysfunction within the tumor microenvironment undermines the control of cancer progression. NK cells play critical roles in limiting early tumor growth and metastatic disease, however, established cancers contain a phenotypically distinct, tumor-specific NK cell compartment. The temporal dynamics, mechanistic underpinning and functional significance of this tumor NK pool remains incompletely understood. To address this, we exploited photo-labeling, combined with longitudinal transcriptomic and cellular analyses, to interrogate the fate of NK cells after tumor entry. In multiple murine cancer models we reveal that conventional NK cells are continuously recruited into tumors, but rapidly adopt a distinct phenotypic state with features associated with tissue-residency and complete loss of effector functions (including chemokine and cytokine production and cytotoxicity), within 48-72 hrs of entering the tumor. Depletion of NK cells from established tumors did not alter tumor growth, indicating that intratumoral NK cells cease to actively contribute to anti-tumor responses. Furthermore, comparable NK populations were identified in human colorectal cancers, confirming translational relevance and raising the possibility that interventions to reactivate NK cells within tissues may boost anti-tumor immunity in established cancers. Indeed, administration of IL-15:IL-15Ra complexes prevented the loss of NK cell function and improved tumor control, generating intratumoral NK cells with both enhanced tissue-residency characteristics and effector function. Collectively, our data reveals the fate of cNK cells after recruitment into tumors and provides insight into how intratumoral NK cell functions may be revived.

**Summary:** Conventional NK cells recruited from the circulation rapidly establish a tissue-resident phenotype defined by impaired cytotoxicity and chemokine production after tumor entry; administration of IL- 15:IL-15Rα complexes further promotes this tissue-residency programme but maintains core NK cell effector functions within the tumor.

## Introduction

The clinical impact of immune checkpoint blockade (ICB) and chimeric antigen receptor (CAR) lymphocytes has prompted extensive research into the composition and function of immune cells within tumors (*1–4*). Amongst these, tumor cytotoxic CD8 T cells have been the most extensively characterized, with a spectrum of exhausted T cells with diminishing effector functions described (*5–8*). Innate lymphocytes, classically represented by Natural Killer (NK) cells, but which now also includes other innate lymphoid cell (ILC) populations, also have potent cytotoxic potential. There is increasing interest in therapeutically harnessing this intratumoral compartment in oncology, however, how these cells behave, or change within the TME remains poorly understood. Unlocking the cytotoxic potential of this expanded pool of effector cells has clear therapeutic promise for cancer patients, particularly in synergizing with ICB.

Conventional NK (cNK) cells are circulatory in mouse and human with a robustly cytotoxic phenotype; they are armed with an array of granzymes and perforin, alongside multiple apoptosis-inducing ligands including TRAIL and FASL (*9–11*). *In vivo*, NK cell deficiency results in impaired control of tumor growth and enhanced metastatic disease in multiple murine cancer models (*10, 12*). Importantly, beyond their direct cytotoxic functions, NK cells further act as orchestrators of the T cell response through their recruitment, activation and expansion of dendritic cells (DCs), particularly conventional type 1 DCs (cDC1), via their production of the chemokines CCL5, XCL1, as well as FLT3L and IFNγ (*11, 13, 14*). The cDC1 subset is the most adept at trafficking to the draining lymph node to cross-present tumor antigens to naïve T cells, and thus drive the expansion of anti-tumor CD8 T cells (*15, 16*). Furthermore, cDC1 within the TME attract and restimulate effector CD8 T cells to amplify and sustain the response (*17, 18*). Collectively, these data emphasize the key role of coordinated NK cell-DC crosstalk for generating durable anti-tumor responses. Indeed, the frequency of NK and cDC1 within melanoma is predictive of the response to anti-PD-1 therapy (*14*).

However, phenotypic heterogeneity and dysfunction has been described within the intratumoral NK cell compartment (*19, 20*), likely undermining control of tumor growth. The discovery of different ILC populations involved in type 1 immune responses, collectively termed ILC1s, adds a further level of complexity to understanding the composition of tumor infiltrating lymphocytes (TILs) and whether populations derive from cells recruited from the circulation or tissue-resident compartments (*21–24*). ILC1s share many phenotypic similarities and transcriptional programs with NK cells and despite most ILCs being tissue-resident, circulating ILC1s have been observed in mice (*25–29*). While NK cells are developmentally distinct to ILC1s (*25, 30, 31*) and were classically further distinguished based upon cytotoxic function, more recent descriptions have revealed that ILC1s express granzymes and are able to kill target cells, blurring current understanding of the distinctions between these cell types (*24, 32, 33*). Thus, the dysfunctional compartment of NK cells described within tumors may reflect the adaptation of cNKs to the TME, the contribution of local ILC populations, or a combination of both. Defining how and why intratumoral NK/ILC1 populations form will facilitate more precise investigation of the mechanisms that need to be targeted to manipulate these innate lymphocytes to promote anti-cancer responses.

Here we used temporal labeling of tumors through photoconversion (*34*) to track the fate of NK cells *in vivo* after recruitment into solid tumors. Our data reveals the rapid loss of chemokine and cytokine production, alongside impaired cytotoxicity, as cNK cells adapt to, or are modulated by, the TME. We demonstrate that all cNK cells retained within the tumor ultimately adopt a distinct, dysfunctional state, characterized by expression of CD49a, and that the heterogeneity observed across many pre-clinical tumor models reflects the time cNK cells have spent within the tumor. Depletion of NK cells from established tumors had no impact on tumor growth indicating that these cells have ceased to actively contribute to tumor control. The loss of NK cell functionality after tumor entry could be blocked through enhanced IL-15 signaling, leading to improved control of tumor growth. Collectively, our data clarify the fate of NK cells within solid tumors, defining the tumor -adapted state that these cells form in response to multiple cues within the TME, curtailing their ability to contribute to the anti-tumor response. These data inform further efforts to revive intratumoral NK cells, in combination with ICB, to enhance anti-tumor immunity.

## RESULTS

### NK cells rapidly alter their transcriptome after tumor entry

Recent studies have highlighted that the intratumoral NK cell compartment is distinct, characterized by altered functions including reduced cytotoxicity and an impaired ability to orchestrate DC recruitment and activation (*19–21, 33*). Precisely why this occurs remains unresolved. To date, *in vivo* studies tracking the fate of NK cells specifically within the tumor are lacking. Using our recently described dynamic labeling of the tumor immune compartment (*34*), we sought to define the state of NK cells as they enter tumors and map real time changes in their phenotype and function over time. To this end, MC38 tumors were grafted subcutaneously on the flank of Kaede photoconvertible mice, a transgenic model in which all cells express a green fluorescent protein that irreversibly switches to a red form upon exposure to violet light (*35*). The entire immune compartment of the tumor was selectively labelled (*34*) at 13 days post-engraftment and tissue harvested 2 days later. Thus, amongst the cells isolated from the tumor, unlabeled Kaede Green+ (KG+) cells had been in the tumor for up to 48 hrs, while Kaede Red+ (KR+) cells had spent at least 48 hrs within the tissue. To capture transcriptomic changes across the TIL compartment over time and in an unbiased manner, we employed scRNA-sequencing (scRNA-seq). TILs were FACS-isolated to better equilibrate numbers, further split into KG+ and KR+ populations and analyzed by droplet-based scRNA-seq (Fig. S1A). The gating strategy used ensured the NK cell compartment was fully captured (Fig. S1B).

After quality control, a total of 46,342 TILs were analyzed, comprised of a small number of B cells, putative NK cells and multiple T cell populations (CD4, CD8, Treg, γδ TCR+) as defined by canonical markers (Fig. S1C-E). We focused upon the putative NK cell cluster of 11,808 cells, initially defined by high expression of *Prf1*, *Ncr1*, *Klrb1c* and *Fcgr3*. Unbiased reclustering of these cells in isolation revealed 8 clusters, of which 6 were defined as NK cells based upon expression of *Ncr1, Eomes, Gzmb* and the absence of *Cd3e* and *Il7ra*, including a cycling NK cell cluster additionally expressing *Mki67* and *Birc5* (Fig. 1A, B, Fig. S1F). The remaining two clusters were identified as NKT cells and an ILC cluster, distinguished from *bona fide* NK cells by expression of *Il7ra, Rora* and *Gata3,* in the absence of *Eomes* and *Cd3e* expression (Fig. S1G-I) (*31, 36*). Expression of *Tbx21*, but not *Eomes*within this small cluster indicated the prevalence of ILC1s (Fig. S1I) (*31*).

**Fig. 1.**
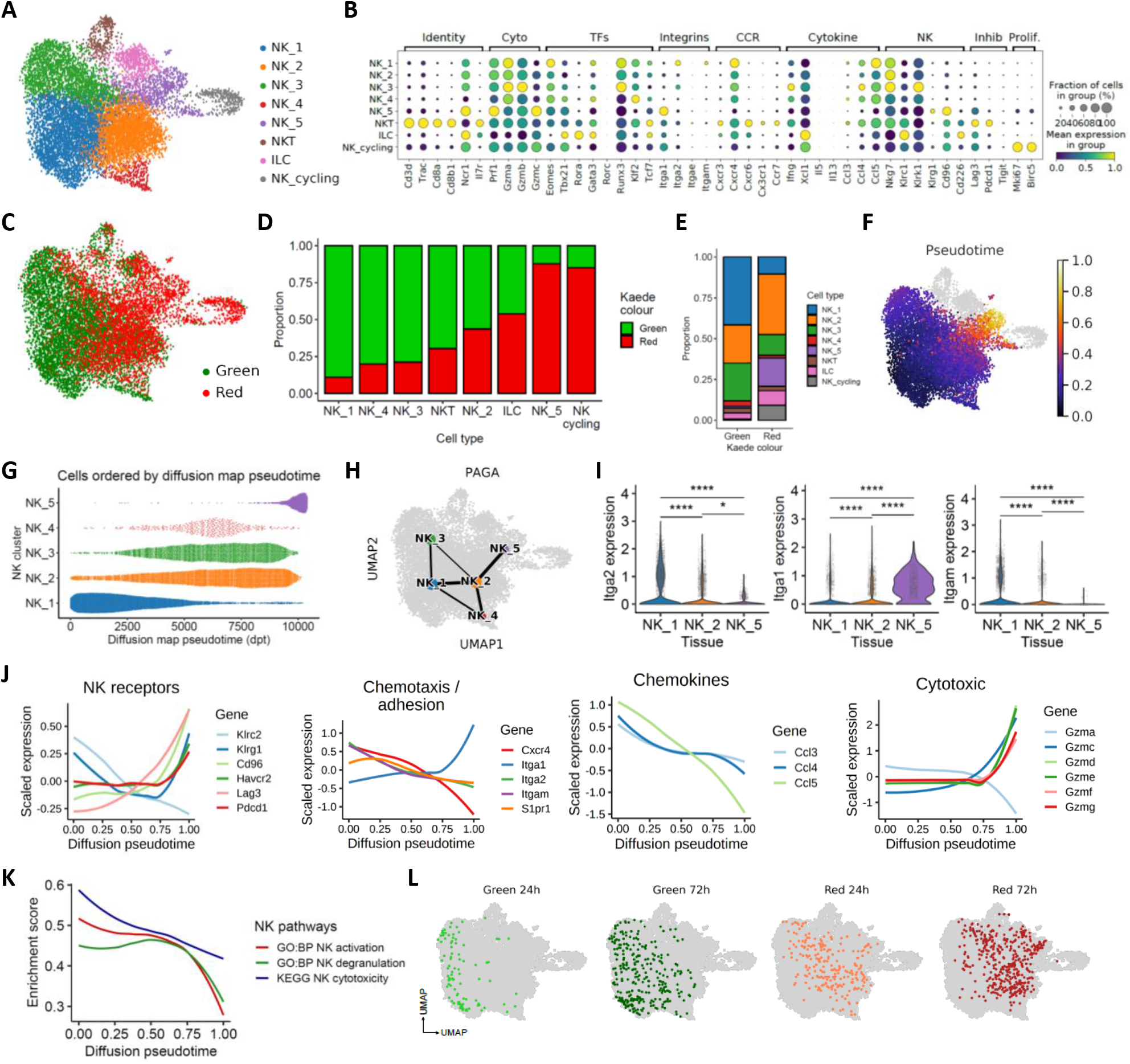
Rapid changes to the NK cell transcriptome after entry into the TME. MC38 tumors, grafted subcutaneously on the flank, were photoconverted and analyzed 48 hrs later using droplet-based scRNA-sequencing. (A) UMAP showing 11808 NK cells defined by expression of *Ncr1*, *Prf1*, *Klrb1c*, *Fcgr3*, resolved into 8 clusters comprised of NK_1 to NK_5, alongside 1 cycling cluster and two further clusters that describe ILC and NKT cells. (B) Dot plots showing expression of selected genes used to further characterize the clusters. (C) UMAP showing the distribution of Kaede Green+ and Kaede Red+ cells across the NK clusters. (D) The proportion of Kaede Green+ and Kaede Red+ cells within each cluster. (E) Proportion of each cluster within the Kaede Green+ and Kaede Red+ cells. (F) UMAP showing diffusion pseudotime trajectory rooted in NK_1. (G) Bee swarm plot of the 5 NK clusters (NK_1 to NK_5) over pseudotime. (H) PAGA illustrating gene expression relationship between NK clusters. (I) Violin plots showing expression of *Iga1*, *Itga2*, *Itgam* across NK_1, NK_2 and NK_5. (J) Differentially expressed genes over pseudotime grouped by function. (K) Pathway analyses characterizing changes in NK activation, degranulation, and cytotoxicity over pseudotime. (L) UMAPs showing integration of NK cell data after 24 and 72 hrs post-photoconversion with the original data derived from 48 hrs post-labeling.

We then determined the proportion of KG+ and KR+ cells within each cluster to contextualize the time spent within the tumor microenvironment of each subcluster (Fig. 1C-E). These data revealed a gradation in the proportion of KR+ cells across the NK clusters, with the NK_1 cluster comprised almost entirely of KG+ cells and the NK_5 cluster at the other extreme, consisting of approximately 90% KR+ cells. The high proportion of KR+ cells within the NK_cycling cluster suggests that some NK cells retained within the tumor are maintained by local proliferation. To temporally order the transcriptomic changes that occur within NK cells after entering the tumor, we performed a pseudotime trajectory analysis, rooted in NK_1 since this contained the highest proportion of NK cells that had recently entered the tumor. These analyses highlighted the NK_5 cluster as the end state of the trajectory (Fig. 1F, G). Similarly, partition-based graph abstraction (PAGA) to map cluster connectivity (Fig. 1H), indicated that the major cell fate trajectory for tumor NK cells progresses from NK_1 through to NK_2 and then onto NK_5 clusters.

Differential expression of the integrins CD49a and CD49b has been used to distinguish cNK cells (CD49b+) and tissue-resident NK cells and ILC1 (CD49a+) (*21*). Expression of *Itga2* (CD49b) was limited to the NK_1 and NK_2 clusters while *Itga1* (CD49a) was expressed by the majority of the NK_5 cluster, but not the other clusters (Fig. 1l). Interestingly, expression of *Itgam*, which encodes CD11b and defines mature NK cells in the circulation (*37*), was limited to the NK_1 cluster. To begin investigating the transcriptomic changes occurring within intratumoral NK cells, we plotted the expression of functionally grouped genes over pseudotime (Fig. 1J) (*38*). As NK cells progressed towards the terminal intratumoral pseudotime state, there was an upregulation of inhibitory receptors including *Pdcd1*, *Lag3*, *Havcr2*, *Cd96*, a change in migration-associated transcripts with a marked reduction in *Cxcr4* and *Itgam* expression, but an increase in *Itga1*, as well as loss of expression of *Ccl3*, *Ccl4*, *Ccl5* (chemokines associated with DC recruitment), and altered granzyme expression.

Consistent with these findings, geneset enrichment analysis identified a reduction in ‘*NK activation’*, ‘*NK degranulation’* and ‘*NK cytotoxicity’* pathway gene sets over pseudotime (Fig. 1K).

To better understand changes in intratumoral NK cells over time, we additionally analyzed NK cells in scRNA-seq data where TILs were assessed at 24 and 72 hrs post photoconversion (*34*), providing finer granularity of the changes that occur over real time. Re-analysis of 1035 NK cells revealed three NK clusters (Fig. S2A, B). Notably, the NK_a cluster was comprised almost entirely of KG+ cells, while NK_c contained only KR+ cells, consistent with a simple linear trajectory (Fig. S2B, C). Substantial changes in gene expression across these three clusters closely matched those observed in our initial analysis, including a switch from *Itgam* to *Itga1* expression (Fig. S2D, E). Finally, to determine where NK cell transcriptomes from 24 and 72 hrs post photoconversion would embed along the pseudo-time trajectory, we projected these data into our initial reference single cell data set (Fig. 1F). Reassuringly, these time-stamped data closely aligned to our pseudotime trajectory, validating that our analyses faithfully model temporal transcriptional changes (Fig. 1L).

Collectively, these data reveal rapid changes to the transcriptome of *Itgam* expressing cNK cells after entering the TME from the circulation. Once within tumors, NK cells rapidly differentiate towards a common transcriptional state characterized by substantially altered core functions and expression of *Itga1*.

### Differential expression of CD11b and CD49a expression capture temporal changes in NK cell phenotype

To validate the rapid changes observed in the transcriptional profile of NK cells entering the tumor, we turned to flow cytometry and initially sought to establish how best to identify cells described along the main cell fate trajectory. Given the differential expression of *Itga1* and *Itga2* (Fig. 1I) and prior use of these integrins to define NK populations (*19*), we assessed CD49a versus CD49b on CD3- NK1.1+ cells in MC38 tumors. However, our analysis revealed only two clear NK cell populations (Fig. 2A), prompting us to try alternative gating strategies. Comparison of CD49a versus CD11b expression identified three populations of CD3- NK1.1+ cells: CD11b+ CD49a- cells, CD11b-CD49a- and CD11b- CD49a+ cells (Fig. 2B). Importantly, a similar staining pattern was observed across multiple murine cancer models including other subcutaneous (CT26, B16F10-OVA), orthotopic (E0771) and primary (PyMT) tumors (Fig. 2C, D). To extend this analysis and compare intratumoral NK cells with the NK/ILC1 populations found in healthy tissue, we assessed the CD3- NK1.1+ cells in MC38 and B16-F10 tumors, alongside spleen, liver, small intestine and colon (Fig. S3). Using a flow cytometry panel (Fig. S3a) that included ‘markers’ associated with NK cells (CD49b, EOMES, CD11b), and ILC1s (CD49a, CD200r1, CXCR6, DNAM-1), these data indicated that the tumor NK compartment was phenotypically distinct to the ILC1 populations identified in non-tumor tissues, consistent with recent scRNA-seq analyses (*21*). While clustering closer to the splenic NK cells, which are dominated by mature cNK, a spectrum of states was clear including CD49a+ cells specific to the tumor.

**Fig. 2.**
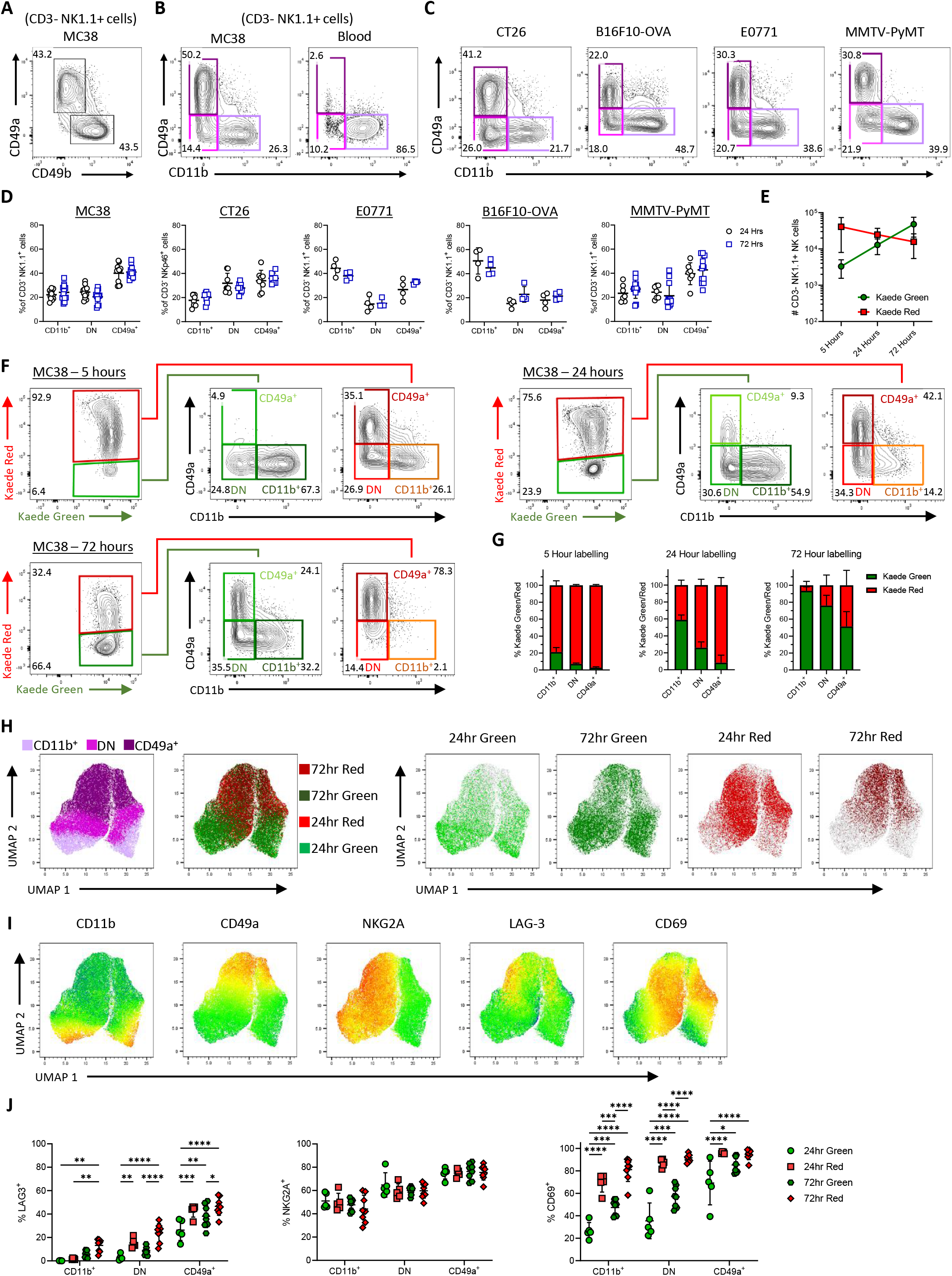
Differential expression of CD11b and CD49a capture temporal changes in NK cell phenotype. (A) Expression of CD49a vs CD49b by NK cells (CD3- NK1.1+) isolated from MC38 tumors grafted subcutaneously on the flank. (B) Expression of CD49a vs CD11b by NK cells isolated from MC38 tumors, alongside blood. (C) Expression of CD49a vs CD11b by NK cells across multiple tumor models. NK cells identified as CD3- NK1.1+ in C57BL/6 mice, CD3- NKp46+ in BALB/c mice. (D) Proportion of NK cells within the CD11b+ CD49a-, CD11b- CD49a- (double negative, DN) and CD11b- CD49a+ gates across the different tumor models. Data collect from B16F10-OVA and EO771 tumors was from 1 independent repeat, and MC38, CT26, and MMTV-PyMT tumors pooled from 2 independent repeats, where MC38 (24 hr n=12, 72hr n=12), CT26 (24 hr n=8, 72hr n=8), EO771 (24 hr n=4, 72hr n=4), B16F10-OVA (24 hr n=4, 72hr n=4), and MMTV-PyMT (24 hr n=7, 72hr n=12). (E) Number of Kaede Green+ and Kaede Red+ NK cells at 5, 24 and 72 hrs post photoconversion of MC38 tumors. (F) Expression of CD49a versus CD11b by Kaede Green+ and Kaede Red+ NK cells at 5, 24 and 72 hrs post photoconversion. (G) The proportion of Kaede Green/Red for each NK cell subset at each time point post photoconversion. Data at 5 (n=5) hrs is representative of 1 independent repeat, whereas 24 (n=8) and 72 (n=11) hrs pooled from 2 independent repeats. (H) UMAPs showing protein expression of CD11b and CD49a alongside Kaede Green/Red expression by NK cells isolated from MC38 tumors at 24 and 72 hrs post photoconversion. (I) UMAPS showing expression of CD11b, CD49a, NKG2A, LAG-3 and CD69. (J) Enumeration of the proportion of cells expressing NKG2A, LAG-3 and CD69 across the NK cell subsets. Data in UMAPs 24hr (n=5), and 72hr(n=8) are representative of two independent repeats. Statistical significance was determined by two-way ANOVA with Šidák’s multiple comparisons test (J). *P<0.05, **P<0.01, ***P<0.001, ****P<0.0001.

To confirm that expression of CD49a versus CD11b could be used to assess the phenotype of NK cells as they enter the tumor and then track changes over time, we photoconverted MC38 tumors and analyzed the intratumoral NK cell compartment 5, 24 and 72 hrs later. Enumeration of the total number of NK cells indicated the continuous recruitment of new (KG+) cells over this time frame alongside a modest decline in the number of retained (KR+) cells within the tumor (Fig. 2E). Analysis at only 5 hrs post-photoconversion revealed a small KG+ population of newly arrived NK cells, the majority of which were CD11b+ CD49a- and the CD49a+ compartment was absent (Fig. 2F). Analysis at 24 hrs post-photoconversion revealed that approximately 10% of the KG+ NK cells expressed CD49a, establishing that a switch in integrin expression begins to occur after only 1 day in the TME. Notably, the KR+ population at 72 hrs post photoconversion, which by definition had spent at least 3 days within the tumor, were essentially all CD49a+ CD11b-. Since a mixture of CD49a+ and CD49a- NK cells were evident amongst the KR+ cells at 24 hrs, these data indicate that full conversion of cNKs takes more than 24 hrs but is completed within 3 days. Considering the proportion of KG vs KR cells of each subset at the different time points highlighted that the % of KG+ cells within each population increased over time, so by 72 hrs post photoconversion, the vast majority of the CD11b+ CD49a- and CD11b- CD49a- populations comprised of cells newly entering since labeling (Fig. 2G). Importantly, comparable dynamic changes in NK cell phenotype over time were observed in other murine tumor models, both across genetic backgrounds and in tumors targeting other anatomical sites (Fig. S4). In addition, we further validated the expression patterns observed in our transcriptomic analysis, identifying increased expression of LAG-3 and CD69 as characteristic features of the CD49a+ ‘tumor - retained’ state (Fig. 2H-J). Interestingly, expression of NKG2A, the inhibitory receptor encoded by *Klrc1*, showed a biphasic expression pattern, largely independent of time within the tumor.

These data clearly indicate that intratumoral NK cell heterogeneity can be explained by the time cNK cells have spent within the tumor; cNK cells enter the tumor from the circulation and it takes approximately 24 hrs for these cells to start upregulating CD49a. The absence of CD49a- NK cells amongst the KR+ NK cells at 72 hrs post-photoconversion indicates that these cells must all either differentiate to a CD49a+ state, egress the tumor, or die *in situ*. To investigate NK cell egress, tumors were photoconverted and the KR+ cells present in the dLN and spleen analyzed 24 and 72 hrs later (Fig. S5). The proportion and total number of KR+ NK cells in either tissue was very low, indicating that few NK cells egressed the tumor. Of those NK cells that did egress, the majority lacked CD49a expression. Collectively these data indicate that the CD49a+ NK cell compartment in these murine tumor models arise from the differentiation and retention of circulating cNK cells.

### NK cells rapidly lose core effector functions after tumor entry

Having established how to identify the temporal changes in NK cell state through a combination of photolabeling and analysis of integrin expression, we investigated alterations to NK cell function over time in the tumor. NK cells orchestrate the accumulation and activation of DCs within the tumor through their production of chemokines including CCL3, CCL4, CCL5 (which all bind CCR1, CCR3 and CCR5) and XCL1 (*13, 39*). While *Ccl3*, *Ccl4* and *Ccl5* expression all significantly declined over pseudotime, *Ccl5* transcripts showed the most dramatic decrease. To validate this, intratumoral NK cells were stimulated *ex vivo* and assessed by intracellular flow cytometry. Analysis of NK cells at 24 hrs post-photoconversion revealed that production of CCL5 was restricted to KG+ CD11b+ CD49a-NK cells, confirming the rapid loss of CCL5 expression after tumor entry (Fig. 3A). NK cell production of IFNγ is a further mechanism by which innate immune cells may activate DCs to enable priming of antigen specific CD8+ T cells (*40*). Post-stimulation *ex vivo*, the majority of KG+ and KR+ intratumoral NK cells failed to produce IFNγ, unlike splenic NK cells under these conditions (Fig. 3B). To further investigate NK cell *Ifng* expression in the absence of *ex vivo* stimulation, we turned to *Ifng*^cre/mKate2^ reporter mice. To confirm accurate reporting in this newly generated model, splenocytes were stimulated *ex vivo* with recombinant IL-12 and IL-18, which resulted in robust mKate2 reporting of *Ifng* expression in NK cells but not T cells (Fig. S6A). However, NK cells freshly isolated from MC38 tumors grafted into *Ifng*^cre/mKate2^ mice, lacked mKate2 expression regardless of their integrin expression (Fig. 3C). To validate the changes in the cytotoxic profile of intratumoral NK cells indicated by the transcriptomic analysis (Fig. 1J), we confirmed Granzyme A was produced by the vast majority of CD11b+ CD49a- cells, but significantly less was detected in the CD11b- CD49a- and CD11b- CD49a+ populations (Fig. 3D). Contrasting with the decline in Granzyme A production, and again consistent with the changes observed over transcriptional pseudotime, Granzyme C production was lacking from KG+ CD11b+ CD49a- cells and most highly produced by KR+ CD11b- CD49a+ cells (Fig. 3E). Granzyme B, which was not differentially expressed over pseudotime at the transcriptional level, was detected across all the NK cell populations and a significantly higher proportion of KR+ NK cells produced Granzyme B (Fig. 3F). However, production of Perforin was significantly reduced in CD11b- CD49a+ NK cells versus the CD11b+ CD49a- populations, and the proportion of NK cells expressing CD107a was significantly reduced amongst the KR+ compartment (Fig. 3G, H).

**Fig. 3.**
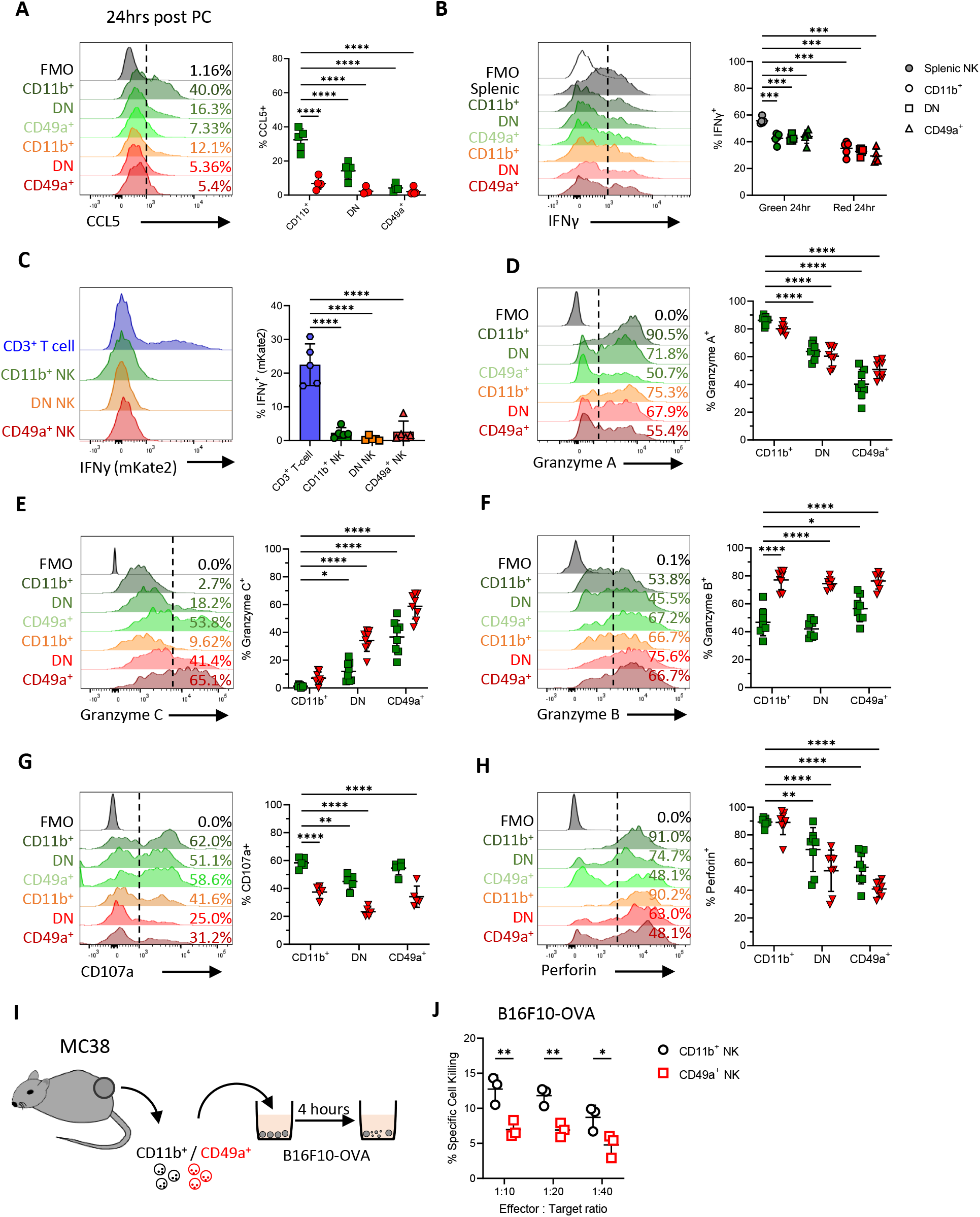
NK cells rapidly change chemokine, cytokine and granzyme production within 24 hrs of entering the TME. Flow cytometry was used to validate changes to the function of NK cells isolated from MC38 tumors at 24 hrs post photoconversion. (A) Proportion of NK cells producing CCL5 after *ex vivo* stimulation, with representative histograms alongside enumeration (n=5). (B) Proportion of NK cells producing IFNγ after *ex vivo* stimulation, with representative histograms alongside enumeration (n=5). (C) Reporting of mKate2 in using Ifnγ^mKate2^ reporter mice, with T cells versus NK cells isolated from MC38 tumors assessed. Representative histograms and the proportion of mKate2+ cells shown. Representative histograms showing proportion of NK cells producing (D) Granzyme A, (E) Granzyme C, (F) Granzyme B, (G) CD107a, (H) Perforin in MC38 tumours that were photoconverted and analysed 24hrs later. Data pooled from 2 independent experiments (n=8) for all analyses except CD107a expression where n=5 from 1 independent experiment. (I) Cartoon showing experimental setup whereby CD11b+ and CD49a+ NK cells from MC38 tumors were FACS-isolated and co-cultured with B16-F10 melanoma cells pre-treated with cell trace violet that were then stained for cell viability. (J) NK-mediated target cell killing comparing cytotoxicity between FACS sorted CD11b+ and CD49a+ NK cells (n=3). Significance was determined by two-way ANOVA with Šidák’s multiple comparison test (A, B, and D – H) comparing means to Kaede Green+ CD11b+ cells, or between groups (J), and a Fruskal- Wallis test with Dunn’s multiple comparison test (C). *P<0.05, **P<0.01, ***P<0.001, ****P<0.0001.

These changes in the function of NK cells when retained within tumors were further confirmed in other murine tumor models (Fig. S6B-E). Of note, having analyzed CT26 tumors grown subcutaneously, we further asked whether dysfunctional NK cells were also observed when this colorectal cell line was grown orthotopically (Fig. S6F, G). Here, in addition to EOMES+ NK cells within the CD3- NKp46+ populations, CXCR6+ EOMES- ILCs were evident, a population lacking in the subcutaneous model that reflected the contribution of ILCs present within the local tissue (Fig. S6H).

The NK cells within orthotopically implanted CT26 could again be divided into three subsets based on CD49a versus CD11b expression, and the CD49a+ NK cells expressed Granzyme C rather than Granzyme A and had reduced perforin expression (Fig. S6I). To directly assess whether cNK cells differentiate into the CD11b- CD49a+ phenotype, we MACS-isolated splenic NK cells from Kaede mice and transferred them i.v. into WT C57BL/6 hosts bearing MC38 tumors (Fig. S7A). Only within the MC38 tumors did the transferred KG+ NK cells upregulate CD49a and Granzyme C (Fig. S7C-G). Finally, we sought to test whether the CD11b-CD49a+ NK population was less cytotoxic that the CD11b+ CD49a- cells that enter tumors. To do this, we FACS-isolated the CD11b+ and CD49a+ NK cells from MC38 tumors and tested their ability to lyse B16F10 tumor cells *in vitro* over 4 hrs (Fig. 3I). The CD11b+ population showed significantly better killing of the labelled CD49a+ NK cells, consistent with functional differences in the cytotoxic ability of these NK cell populations (Fig. 3J). Collectively, these data demonstrate the presence of dysfunctional NK cells across multiple preclinical cancer models targeting different tissues. In all, upregulation of CD49a expression in the absence of CD11b is associated with retention in the TME, altered effector functions and less efficient tumor cell killing.

### Multiple mechanisms in the TME drive the conversion of cNK cells to a tumor - retained CD49a+ NK cell compartment

Having defined the cellular state adopted by cNK cells that become retained within tumors, we sought to understand the mechanisms within the TME that drive this cell fate. The TGFβ and PGE_2_ pathways have both been linked to altered NK cell functions both *in vitro* and *in vivo* (*13, 19, 20*). We initially asked whether NK cells with the phenotype and functional repertoire of CD49a+ ‘tumor - retained’ cells, could be induced by exposure of cNKs to recombinant TGFβ and/or PGE_2_. Culture with TGFβ but not PGE_2_ caused the upregulation of CD49a, however CD11b expression did not change (Fig. S8A). Culture with either TGFβ or PGE_2,_ alone, or in combination, diminished NK cell production of CCL5 upon restimulation, while complete loss of IFNγ expression required both TGFβ and PGE_2_ exposure (Fig. 4A, Fig. S8B). However, no alteration in Granzyme A or Granzyme C production was observed, indicating that the full spectrum of changes associated with the NK cells retained within tumors could not be recapitulated *in vitro*. We further repeated our *in vitro* cultures at 1% oxygen to better model potential conditions in the TME and the potential response to hypoxia in altering NK cell function (*41*). In addition, only very modest changes in integrin and granzyme expression were observed when cultures under normoxia versus hypoxia were compared (Fig. S8C-G).

**Fig. 4.**
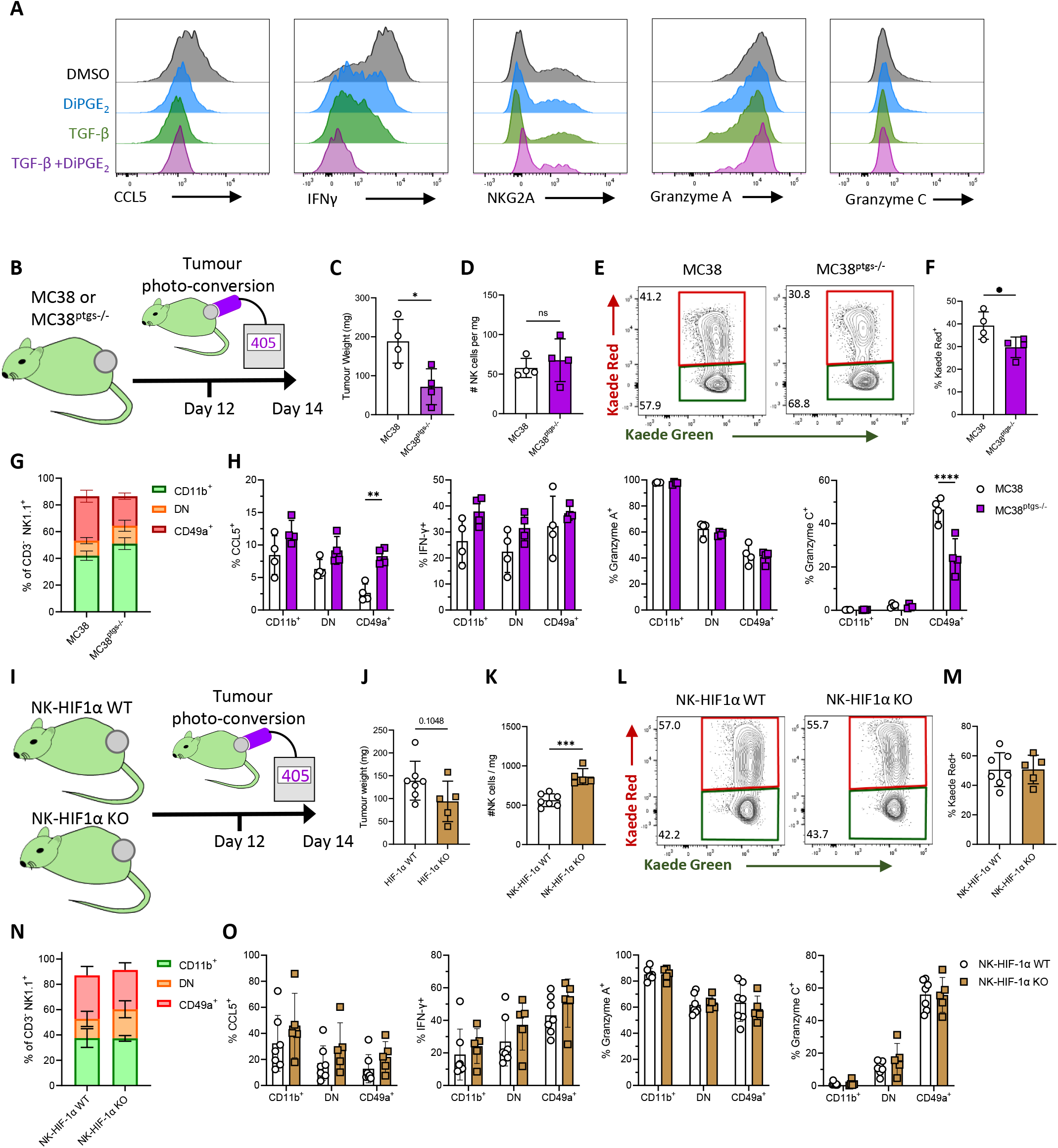
Multiple mechanisms in the TME drive the conversion of cNK cells to a tumor-retained CD49a+ state. The role of TGFβ, PGE2 and Hif1a were investigated *in vitro* and *in vivo* as mechanisms promoting NK cell differentiation to the tumor retained state characterized by CD49a expression and disrupted core functions. (A) Bar chart showing proportion of CD11b+ and CD49a+ NK cells after 48 hrs in culture in RPMI with IL-2/IL-15 further supplemented with TGF-β and/or DiPGE2. (B) Cartoon summarizing experimental design for impeding tumor produced PGE_2_. Kaede mice were grafted with either MC38 (n=4) or MC38^ptgs1-/-ptgs2-/-^ (MC38^ptgs-/-^, n=4) cells, photoconverted on D12 and analyzed 48 hrs later. (C) Tumor weight. (D) Enumeration of NK cells per mg tumour. (E) Flow cytometry plots showing Kaede Green versus Kaede Red expression by NK cells. (F) Proportion of NK cells expressing Kaede Red label 48 hrs post photoconversion. (G) Proportion of NK cells in CD11b+ CD49a-, CD11b- CD49a- and CD11b- CD49a+ subsets. (H) Proportion of NK cells producing CCL5, IFNγ, Granzyme A, Granzyme C after *ex vivo* restimulation. (I) Cartoon summarizing experimental design for targeting Hif1a within NK cells, using Ncr1^cre^ x Hif1a^f/f^ x Kaede mice (NK-HIF1α KO n=5) versus cre-negative littermates (NK-HIF1α WT n=7) grafted with MC38 tumors, photoconverted on D12 and analyzed 48 hrs later. (J) Tumor weight. (K) Enumeration of NK cells per mg tumour. (L) Flow cytometry plots showing Kaede Green versus Kaede Red expression by NK cells. (M) Proportion of NK cells expressing Kaede Red label 48 hrs post photoconversion. (N) Proportion of NK cells in CD11b+ CD49a-, CD11b- CD49a- and CD11b- CD49a+ subsets. (O) Proportion of NK cells producing CCL5, IFNγ, Granzyme A, Granzyme C after *ex vivo* restimulation. Statistical significance was determined by unpaired t-tests (C, D, F, J, K, M) or two-way ANOVA with Šidák’s multiple comparison test comparing group means (H and O). *P<0.05, **P<0.01, ***P<0.001, ****P<0.0001.

While we could partially recapitulate NK phenotype and functional changes *in vitro,* to further define the cues that modulate intratumoral NK cells we turned to *in vivo* approaches, reasoning that these would better model the complex TME. To investigate the role of TGFβ in driving changes to NK cells within the tumor, we sought to deplete intratumoral Tregs, given these are a critical source of TGFβ (*42*). Since NK cells can express CTLA4 and CD25 (*43, 44*), we utilized anti-OX40 Abs, which efficiently deleted the FoxP3+ CD4 T cells within the tumor while leaving NK cells intact (Fig. S8H-J). However, depletion of intratumoral Tregs did not alter the proportions of NK cells differentially expressing CD49a or CD11b (Fig. S8K). No differences in IFNγ, Granzyme A or Granzyme C production were detected, although CCL5 production was significantly enhanced by Treg-depletion (Fig. S8L).

To investigate the impact of tumor cell-derived PGE_2_, MC38 cells deficient COX-1 and COX-2 (encoded by *Ptgs1*/Ptgs2, termed MC38^ptgs-/-^) were grafted on the flank of C57BL/6 Kaede mice and the NK cell compartment compared with that of MC38 tumors (Fig. 4B). Despite impaired tumor growth of the MC38^ptgs-/-^ cells (Fig. 4C), the proportion of NK cells was not significantly altered, and the retention of NK cells only modestly impacted (Fig. 4D-F). While the proportion of the NK cell subsets defined by CD49a and CD11b expression were unchanged, significant differences in CCL5 and Granzyme C production were observed, indicating a partial effect on NK cell phenotype and function (Fig. 4G, H). To specifically test the role of HIF1α in restraining NK cell functions within the tumor we generated Ncr1^cre^ x Hif1a^f/f^ x Kaede mice and grafted these, alongside cre-negative littermate controls with MC38 tumors (Fig. 4I). Tumors were photoconverted on day 12 and analyzed 48 hrs later after *ex vivo* stimulation. Conditional deletion of HIF1α expression in NK cells failed to significantly impair tumor growth (Fig. 4J) and while there was a modest increase in the number of NK cells in the tumor (Fig. 4K), no differences in the % of KR+ NK cells, the proportion of NK cell subsets, nor production of CCL5, IFNγ, Granzyme A or Granzyme C was detected (Fig. 4L-O). Collectively, these data recapitulate published observations regarding the role of TGFβ and PGE_2_ in altering NK cell function, particularly CCL5 production. However, numerous signals, rather than a single mechanism appear to cause the full array of changes characteristic of NK cells that become retained within murine tumor models.

### NK cells in human colorectal cancer show loss of effector functions

Having demonstrated across multiple murine cancer models that the NK cells retained within tumors become dysfunctional, we sought evidence that NK cells undergo a comparable transition in human cancers. NK cells are present in colorectal cancer, but most adopt a “resting”, non-activated state (Fig. S9A, B) (*45*). To precisely characterize their phenotype, we re-analyzed scRNA-seq of human colorectal tumors (*46*), including 3521 NK cells from 62 patients (Fig. 5A, S9C, D). Among these, the majority were *FCGR3A*-expressing, *SELL*-negative, *CD3D*-negative NK1 cells, which are the CD56^dim^CD16^+^ cytotoxic NK cell subset in humans (*47*) (Fig. 5B). Within NK1 cells, unbiased clustering revealed 3 distinct cell states, including an *ITGAM*-expressing NK1-1 subset, an *ITGA1*-expressing NK1- 3 subset, and a double-negative NK1-2 subset, similar to our observations in murine tumors (Fig. 5C). Expression of several granzymes (*GZMB*, *GZMH*, *GZMM*), perforin (*PRF1*), and gene signatures representing NK cytotoxicity and activation were enriched on NK1-1 but downregulated in NK1-2 and NK1-3 (Fig. 5D, E, S9E). This is consistent with our observations in mice, where resident CD49a^+^ cells rapidly downregulate cytotoxicity and activation markers compared to newly-infiltrating CD11b^+^ cells. Moreover, there was reduced expression of *PRF1* and NK cytotoxicity genes in tumor-infiltrating NK cells versus adjacent normal colorectal tissue, further suggesting a loss of anti-tumor NK cell function (Fig. 5D, S9F). The *ITGAM*-expressing subset more closely associated with early-stage T1-3 tumors, while the *ITGA1*-expressing more densely populated advanced-stage T4 tumors or tumors that had progressed to disseminate lymph node metastases (Fig. 5F, G). Thus transcriptionally, a spectrum of NK cell states, comparable to that observed in murine tumor models was evident in human CRC. To validate these phenotypes at the protein level, we compared NK cells in peripheral blood, with those obtained from unaffected colon and primary CRC. Analysis of paired tissues from 10 CRC patients demonstrated a clear shift from predominantly CD11b^+^ NK cells in blood to more CD49a^+^ NK cells in the tumors, while normal colon was dominated by NK cells expressing neither CD11b nor CD49a (Fig. 5H-J, Fig. S10). Deeper characterization of the phenotype of NK cell subsets within the tumor showed a lower expression of the cytotoxicity-mediating molecules Granzyme B and Perforin as well as CD16 within the CD49a^+^ subset (Fig. 5K, L). This was paralleled by higher expression of NKG2A and the tissue-residency markers CD103 and CD69 on CD49a^+^ NK cells. Collectively these data demonstrate the presence of a CD49a^+^ subset of NK cells with features of tissue residency that lacks the expression of cytotoxic proteins within the CRC TME; confirming the translational relevance of our murine model as a tool to identify and validate therapeutic targets.

**Fig. 5.**
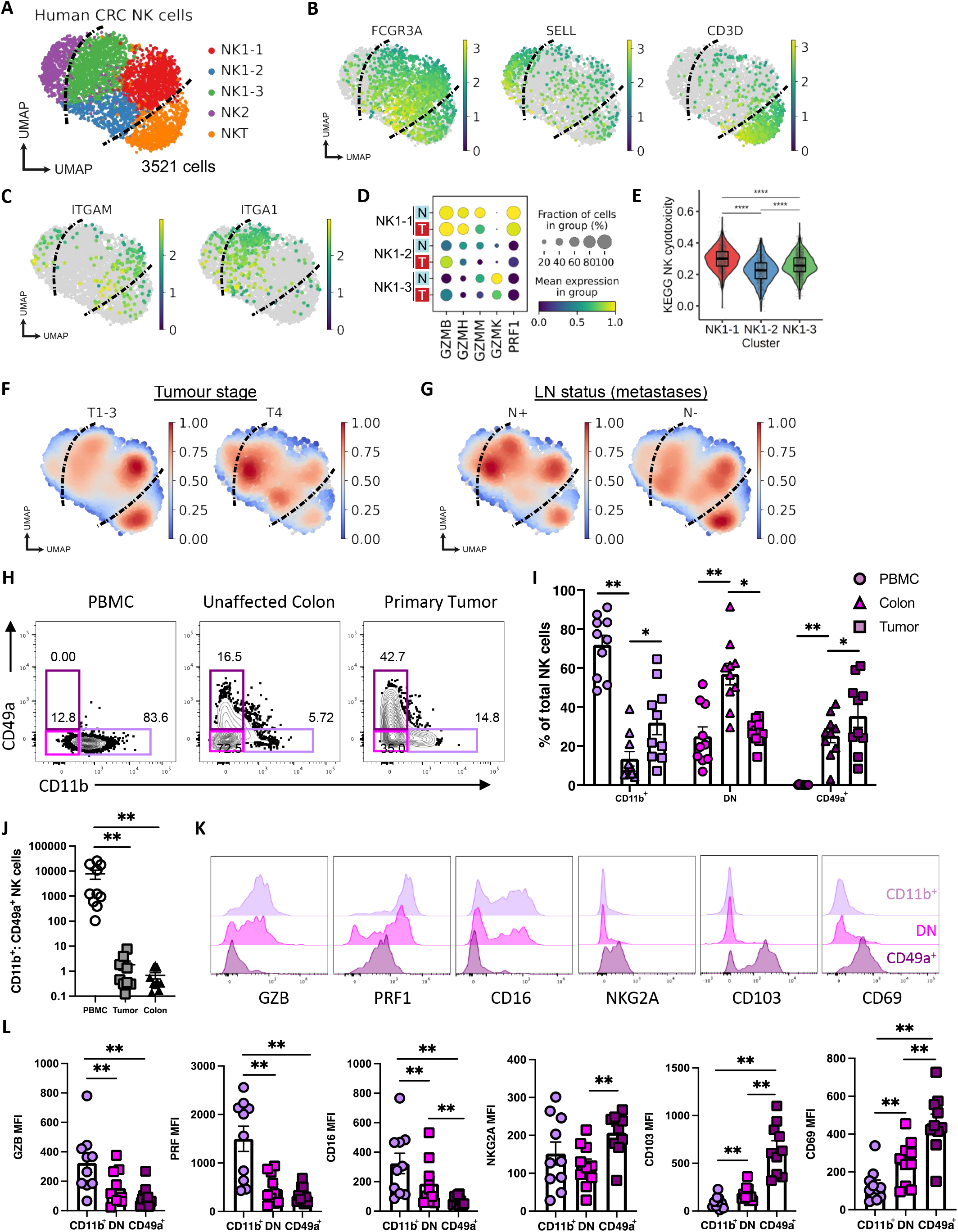
NK cells in human CRC show loss of effector functions. Evidence for dysfunctional NK cells within human CRC was sought through bioinformatics analysis of publicly available data sets alongside flowcytometric analysis of primary human CRC samples. (A) UMAP of 3521 NK cells from scRNA-seq of 62 human CRC samples, from GSE178341. (B) Expression of selected marker genes for clusters shown in ‘A’. (C) Expression of *ITGAM* and *ITGA1* in NK1 subsets. (D) Granzyme and perforin gene expression in NK1 subsets, in tumor (T) versus normal adjacent (N) tissue. (E) Gene set enrichment for KEGG Natural killer cell-mediated cytotoxicity between NK1 clusters. (F) Gaussian kernel density embedding of cells in CRC tumors by tumor staging, and (G) lymph node (LN) metastases. (H) Representation flow plots depicting the frequency of CD49a+ and CD11b+ NK cells across PBMC, unaffected colon tissue and CRC primary tumor for the same donor (n=10). (I) Bar plot showing the frequency of CD11b+CD49a-, CD11b-CD49a- and CD11b-CD49a+ NK cells across compartments. Significance was determined using Wilcoxon test to show specific comparisons. (J) Ratio of CD11b+CD49a- to CD11b-CD49a+ NK cells across tissue compartments. Significance was determined using Kruskal-Wallis. (K) Representative histograms showing expression of selected markers on tumor infiltrating NK cell subsets. (L) Bar plots depicting geoMFI values for specific markers in tumor infiltrating NK cells. Significance was determined using the Kruskal-Wallis test.*P<0.05, **P<0.01, ***P<0.001, ****P<0.0001.

### Enhanced IL-15 signaling drives formation of a distinct cytotoxic intratumoral NK cell population

Our analyses above demonstrate that cNK cells recruited into the tumor rapidly become dysfunctional, both in their cytotoxicity and their ability to recruit and activate DC in the tumor. Furthermore, our data shows that this altered state is the fate of all cNK cells retained within the tumor, questioning the role of these cells in the anti-tumor response. Depletion of NK cells prior to establishing lung metastases resulted in a significant increase in disease burden (*10, 48*). Given their transcriptional and functional state, we hypothesized that within established tumors the intratumoral NK cell compartment is no longer able to contribute to the control of tumor growth. To test this, MC38 tumors were established in WT mice and then NK cells were depleted using anti-NK1.1 Abs, administered from 6 days of tumor growth. Highly efficient NK cell depletion was achieved within the tumor, however, the loss of intratumoral NK cells did not alter tumor growth or mass upon tissue harvest (Fig. S11A-D). To confirm these observations, we repeated the depletion of NK cells within established B16F10-OVA tumors, which have reduced MHCI expression compared to MC38 (Fig. S11E). We grafted tumors into both WT and Rag^-/-^ (C57BL/6 background) mice, the latter to further focus on

NK cell-specific contributions, and again depleted NK cells. While tumors grew much quicker in Rag^-/-^ hosts, depletion of the NK cells from established tumors did not significantly impact tumor growth or mass (Fig. S11F-G). Loss of NK cells in the tumor also resulted in a significant decrease in intratumoral DC numbers (Fig. S11H-K). Collectively these data demonstrate that depletion of NK cells in established tumors does not cause an increase in tumor growth, consistent with the conclusion that intratumoral NK cells fail to meaningfully contribute to the anti-tumor response due to the rapidly acquired dysfunctional state.

Thus, we next sought approaches that could block or reverse the loss of NK cell function after tumor entry to support enhanced anti-tumor responses. IL-15 drives the differentiation and activation of both T and NK cells (*49*). Therapies utilizing IL-15R agonists (ALT-803 and NIZ985) are currently being trialed alone or in combination with ICB (NCT02523469, NCT02452268, NCT03228667, NCT03520686, NCT05096663) and have shown some initial promise in patients that have previously relapsed from immune checkpoint inhibitor treatment (*50*). Initial patient data suggest that the response to IL-15R agonists is associated with NK cell and CD8^+^ T cell expansion and increased IFNγ production. However, it is unclear how IL-15R agonism enhances NK anti-tumor responses or the potential recruitment of NK cells into solid tumors. To determine whether enhanced IL-15 signaling resulted in the maintenance of NK functions normally disrupted by adaptation to the TME, tumor bearing mice were treated with complexes of IL-15:IL-15Ra, since these complexes have a significantly enhanced activity versus recombinant IL-15 alone. We tested these complexes, administered at D7 and D12 of tumor growth, in multiple tumor models (Fig. 6A). BALB/c Kaede mice bearing CT26 tumors showed significantly enhanced tumor control with treatment (Fig. 6B). Analysis of the intratumoral NK cell compartment revealed striking differences in integrin expression following administration of IL-15:IL- 15Ra complexes (Fig. 6C). Treatment resulted in the majority of the NK cells expressing CD49a, and the emergence of a CD49a+ CD11b+ population barely detectable in the PBS controls (Fig. 6D). To ask whether IL-15:IL-15Ra complexes caused the emergence of CD49a+ CD11b+ cells after entry into the tumor, or if these cells were formed in the periphery and trafficked into the tumor, we photoconverted tumors and compared the % KR expression within different intratumoral NK cell subsets. Treatment did not cause a significant difference in the total proportion of KG+ NK cells in the tumor (Fig. 6E). Analysis of the NK cell subsets defined by CD49a versus CD11b expression revealed that the vast majority of CD49a+ CD11b+ NK cells were KR+, demonstrating that this subset emerged over time in the tumor, rather than arising from cells trafficking into the tumor (Fig. 6F). Analysis of NK cell functions revealed that the expanded CD49a+ CD11b+ population observed after IL-15:IL-15Ra treatment had the highest expression of perforin and granzymes A, B, and C (Fig. 6G). Enhanced IL-15 signaling also significantly increased CCL5 production alongside NKG2A expression. Comparable data was observed in both MC38 and B16F10-OVA tumors after IL-15:IL-15Ra treatment, confirming that the CD49a+ CD11b+ NK population that emerged in tumors was not model dependent (Fig. S12). Finally, since enhanced IL-15 signaling impacts both CD8 T cells and NK cells, we sought to determine the NK cell contribution to the enhanced control of tumor growth after treatment. BALB/c Kaede mice bearing CT26 tumors were split into PBS or IL15:IL-15Ra treatment groups, and then further divided into groups receiving isotype control or anti-CD8α Abs to specifically deplete the CD8 T cell compartment (Fig. 6H). Administration of IL-15:IL-15Ra complexes combined with isotype control Abs mediated the greatest control of tumor growth, while CD8 T cell depletion resulted in faster tumor growth than in PBS control mice given isotype Abs (Fig. 6I). Depletion of CD8 T cells in combination with IL-15:IL-15Ra complexes improved tumor control versus PBS/anti-CD8α Abs, consistent with a distinct NK cell and CD8 T cell contribution to anti-tumor responses. Effective depletion of CD8 T cells was confirmed by analysis of IFNγ producing T cells within the tumor (Fig. 6J).

**Fig. 6.**
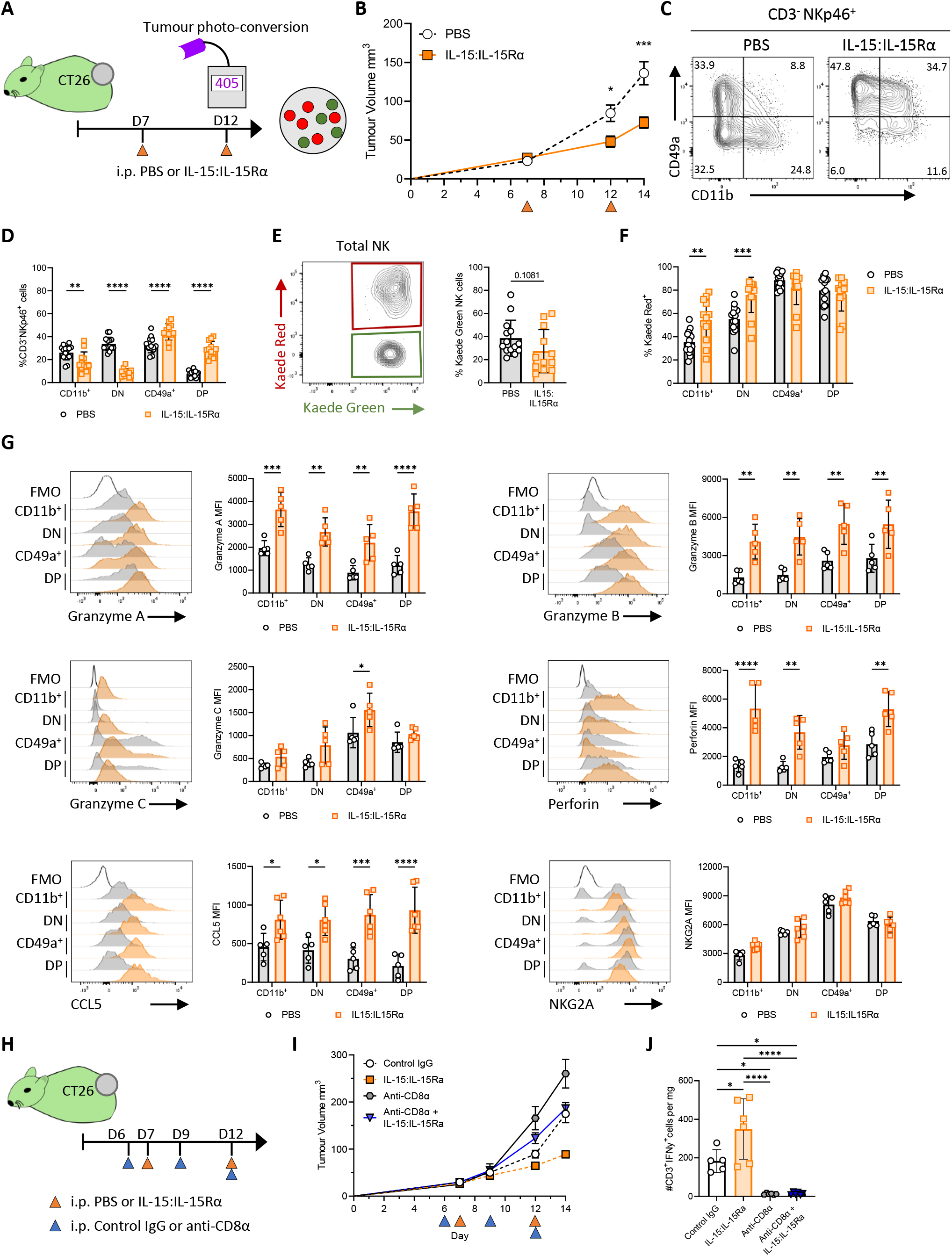
Administration of IL15:IL15Rα complexes result in enhanced tumor control and the formation of CD49+ CD11b+ intratumoral NK cells with heightened functions. To block loss of NK cell functions within the tumor, BALB/c Kaede mice bearing CT26 tumors were treated with IL-15:IL-15Ra complexes. (A) Cartoon showing experimental design. (B) Growth curves for CT26. (C) Representative flow cytometry plots expression of CD49a versus CD11b by NK cells (CD3- NKp46+). (D) Proportion of NK cells with CD11b+ CD49a-, CD11b- CD49a-, CD11b- CD49a+, CD11b+ CD49a+ phenotype. (E) Representative flow cytometry plot showing Kaede Green versus Kaede Red expression by total NK cells alongside enumeration of % Kaede Green+ NK cells. (F) Proportion of Kaede Red+ cells within CD11b+ CD49a-, CD11b- CD49a-, CD11b- CD49a+, CD11b+ CD49a+ subsets of NK cells. Data pooled from 3 independent experiments, (cumulative totals: PBS n=14, IL15:IL-15Ra n=12). (G) Representative histograms and enumeration of Granzyme A, Granzyme B, Granzyme C, Perforin, CCL5, and NKG2A (MFI) after *ex vivo* stimulation (data from 1 of 3 experiments shown, PBS n=5 and IL-15:IL-15Rα n=5). (H) Cartoon showing experimental design for CD8 T cell depletion in combination with IL-15:IL-15Ra complexes (PBS + IgG n=5, PBS + anti-CD8α n=6, IL-15:IL-15Rα + IgG n=6, 15:IL-15Rα + anti-CD8α n=6). (I) Tumor growth curve for experiment illustrated in ‘H’. (J) Total number of NK cells per mg tumor. Statistical significance was determined by two-way ANOVA with Šidák’s multiple comparisons test (B, D, F, G, and I), unpaired t-test (E), or one-way ANOVA with Tukey’s multiple comparisons test (J). *P<0.05, **P<0.01, ***P<0.001, ****P<0.0001.

Combined these data reveal that enhanced IL-15 signaling is able to block the loss of core NK cell functions that occurs as cNK cells adapt to the TME. Administration of IL-15:IL-15Ra complexes results in the emergence of a distinct CD49a+ CD11b+ NK cell compartment, not normally present within tumors, that retains a highly cytotoxic profile and contribute to improved tumor control.

## Discussion

Here we have exploited the site-specific temporal labeling afforded by Kaede photoconvertible mice to determine how NK cells change over time having entered tumours. Multiple studies have described the presence of a distinct CD49a+ NK cell population within murine and human cancers (*19–21, 51, 52*). Our data establishes that this population can be rapidly formed from cNK cells recruited from the circulation that then respond to the TME, and develop a tumour-retained state distinct from that of circulating cNK cells or ILC1s within lymphoid or non-lymphoid tissues. Adaptation to the TME bears hallmarks of tissue-residency, including CD49a and CD69 expression, alongside the loss of the basic cNK effector functions of efficient cancer-cell killing and the recruitment and activation of DCs. This renders intratumoral NK cells inactive in the response, evidenced by their depletion from established tumours failing to impact tumour growth. Having defined the end-state of NK cell differentiation within tumours, we sought to determine the mechanisms that drive these changes. Manipulation of TGFβ, PGE_2_ or the transcription programme controlled by HIF1α, was insufficient to drive the full array of changes that define tumour-retained NK cells, implying that the coordinated action of multiple signals orchestrates this fate. The loss of NK cell functions within the tumour could be prevented through enhanced IL-15 signalling, which pushed intratumoral NK cell differentiation towards a hybrid state defined by high CD49a expression, but also enhanced effector functions. Collectively these data provide new insight into how NK cells change upon recruitment into tumours, the rate at which this occurs, and what this means for the innate lymphocyte compartment in cancers.

The identification of different and potentially tissue-specific ILC1 populations has raised questions as to how the CD49a+ ‘NK cell-like’ compartment of tumours arises and is then sustained over time (*20–22, 33*). This matters for investigations into the reinvigoration of intratumoral NK cells, as well as for the basic understanding of the pre-clinical tools used to develop therapeutic approaches for cancer patients. Our data demonstrates that in multiple commonly used tumour models, the CD49a+ NK compartment is derived almost exclusively from cNK cells. Through tracking real time changes in NK cells after tumour entry, we show that the majority of newly entering NK cells are mature CD11b+ cNK cells, although less mature CD11b- cells were also observed (*37*). Expression of CD49a was absent on this newly recruited population but was induced within 24 hrs of entering the TME. While there is the limited egress of NK cells from the tumour, the majority of these cells lacked CD49a expression, and all NK cells retained within the tumour over several days expressed this integrin. Although the photo-labeling of cells used here is too short to fully assess tissue-residency, of the few NK cells found to egress tumours, the majority lacked CD49a expression. CD49a+ intratumoral NK cells also upregulated CD69, consistent with becoming resident within the tumour, and multiple studies in both mouse and human have previously linked CD49a expression with tissue-residency (*53–55*). Transcriptomically, this population remains distinct from *bona fide* ILC. Using unbiased scRNA-seq analyses, we were able to identify a small, but distinct cluster of ILCs that were transcriptomically distinct from all the NK cell clusters, including those expressing *Itga1*, based upon *Il7ra* and *Rora* expression. Thus, cNK cells establish a ‘tumour-retained’ state that can be partly defined by CD49a expression, rather than ‘converting’ into ILC1s (*19*). However, we struggled to identify a clear CD127+ ILC population by flow cytometry in any of the syngeneic cell line tumour models, limiting further investigation. The lack of CD127+ ILCs within syngeneic cell line tumours likely reflects the basic mechanics of these models, where tumours cells are propagated within a ‘space’ under the skin, rather than within a specific tissue with an associated ILC compartment. Recent studies have explored the origin of intratumoral CD49a+ NK cells using fate-mapping of *S1pr5* expression, an elegant system for investigating the progeny of cNK cells (*33*). Here the authors concluded that most intratumoral CD49a+ innate lymphocytes were not derived from cNK cells based upon S1pr5 fate-mapping, rather, they were a distinct ILC1 population with a history of *Gzmc* expression. This observation is derived from studies using the spontaneous MMTV-PyMT breast tumour model (*56*), with the presence of ILCs within the mammary fat pad potentially explaining the difference with tumour models using cell line implantation. In our hands, ILCs were largely absent from CT26 tumours grown subcutaneously, but a CXCR6+ EOMES- ILC population that expressed Granzyme C was evident within orthotopic CT26 tumours. We observed a comparable CXCR6+ EOMES- NK population in spontaneous MMTV-PyMT tumours. Further studies using fate-mapping of *S1pr5* and *Gzmc,* alongside other approaches to distinguish NK and ILC1 such as *Rora* reporting (*36*), should be employed across the multiple murine cancer models to further resolve contribution of differentiating cNKs cells versus tissue-residents ILCs. While resolving the origin of the CD49a+ NK/ILC1 populations in human cancers is particularly challenging, it is evident that NK or ‘NK-like’ populations characterised by CD49a expression and reduced cytotoxicity are present. In hepatocellular carcinoma these correlated with poor prognosis (*51*).

Particularly striking within our data, was the speed at which cNK cells entering the tumour lost core effector functions. While cNK cells were constantly recruited, 24 hrs within the growing tumour was sufficient for extensive transcriptomic re-programming. By establishing the rate at which NK cells change, our data indicates that enhancing the contribution of innate effector cells within the tumours will be dependent upon over-riding mechanisms operating within the TME, rather than simply enhancing recruitment. Our data further suggests that cellular therapies such as CAR NK cells may be less efficient than hoped in solid cancers if efforts to over-ride tissue-adaptation mechanisms are not appropriately considered and approaches to over-ride the normal differentiation process enacted. A fundamental question not addressed within our study is whether the conversion of cNK to a tumour-retained state is reversible. The extent to which epigenetic changes underpin the transition of cNK cells over time within the tumour, alongside transfer experiments will help to clarify this.

As highlighted above, the changes observed within NK cells over time in the tumour bear hall marks of a tissue-resident programme. Local signalling by TGFβ is well established in driving tissue-residency in both NK cells and CD8 T cells (*19, 57–59*). Our data indicates that local TGFβ signalling within the tumour is insufficient to account for all the changes observed within the retained NK cell compartment, including the clear switch in granzyme production. It was further noteworthy that within the models employed here, no role for the transcriptional programmes controlled by HIF1α was evident in restraining intratumoral NK cell functions, indicating that these pathways may be mouse tumour model specific (*41, 60*). NK cell chemokine and cytokine production appear particularly sensitive to change under the influence of signals within the TME. CCL5 production was lost *in vitro* in the presence of either recombinant TGFβ or PGE_2_, and was the one function that could be enhanced *in vivo* upon targeting either intratumoral Tregs or tumour cell derived PGE_2_. While cNK cells are potent producers of IFNγ, newly developed *Ifng* reporter mice revealed that intratumoral NK cells, regardless of their integrin expression, express very little of this key effector cytokine, again consistent with the rapid loss of this function once within the tumour. Since a defining feature of tissue-resident ILCs is their robust production of effector cytokines, the extent to which the ILC1 compartment really produce IFNγ in orthotopic and spontaneous tumour models should be explored without reliance on restimulation assays.

Having defined the immunological cost of NK cells adapting to the TME, we sought to identify interventions that could limit or block the loss of key functions. IL-15 promotes NK cell survival, proliferation, and cytotoxicity, and therapeutically enhanced IL-15 signalling, alone or in combination with other targets such as PD-L1, is currently being trialled (*50, 61–63*). While recombinant IL-15 failed to impact tumour growth in our hands, IL-15:IL-15Ra complexes which drive superior *in vivo* signalling (*64*), were able to consistently curtail tumour growth in multiple cancer models. Common to all the treated tumours was the upregulation of CD49a expression on intratumoral NK cells, enhanced CCL5 production, and increased granzyme and perforin expression consistent with heightened cytotoxicity and their contribution to the enhanced anti-tumour response. Depletion of CD8 T cells within these tumours indicated a contribution from the revived NK compartment alongside a heightened CD8 T cell response. While systemic administration of IL-15:IL-15Ra complexes did drive splenomegaly, due to peripheral expansion of peripheral NK and CD8 T cell populations (*64, 65*), our data indicates that treatment promoted both a tissue-resident phenotype and superior effector function for NK cells specifically within the tumour. Previous studies have identified that the combination of IL-15 and TGFβ signalling synergised in driving a tissue-resident phenotype defined by CD49a, CD69 and CD103 (*66*).

Our data highlights that the strong activation driven by IL-15:IL-15Ra complexes sustains heightened effector functions despite seemingly promoting aspects of a tissue-residency programme.

In summary, here we have provided a detailed map of how circulating NK cells respond to the TME and their fate within this tissue. These studies highlight how quickly immune cells can become dysregulated within the TME and can support the design of therapeutic approaches designed to revive the innate arm of the anti-tumour response and enhance tumour control.

## Materials and Methods

### Study Design

The main aims of this study were to understand how the NK cell compartment of tumors was formed, determine the mechanisms controlling the fate of NK cells within the tumor and to identify approaches to manipulate NK fate and function within the tumor. We sought to define the fate of NK cells entering the tumor from the circulation through site-specific temporal labeling of the entire immune compartment of tumors. This was achieved using syngeneic tumor cell lines grafted into transgenic mice expressing green photoconvertible proteins which, when exposed to violet light, switched to a red fluorescence that could then be detected for a number of days post photoconversion. Tumors were colonized by the host immune cells (which were ‘green’) and by labeling all the immune cells within the tumor at a given moment in time (i.e. turning them ‘red’), we could distinguish retained and newly entering cells to then unpick the fate of cells over time in the tumor. We use time course experiments and a combination of single-cell RNA-sequencing, to unbiasedly capture transcriptomic changes, followed by flow cytometry to validate these changes at the protein level. Building from this characterization, we then investigated the mechanisms causing the changes in NK cell phenotype and function as well as interventions that could alter the changes that occurred in NK cells after entering the tumor.

### Mice

Female C57BL/6 and Rag2^-/-^ mice 6 – 8 weeks were purchased from Charles River and allowed to acclimate to the University of Birmingham Biomedical Services Unit for a week. C57BL/6 Kaede and BALB/c Kaede mice are maintained and bred at the University of Birmingham Biomedical Services Unit. Hif1a^fl/fl^NCR1^iCreTg^ (kindly provided by Dr. Christian Stockmann, University of Zurich, Switzerland) were crossed with C57BL/6 Kaede mice and maintained at the University of Birmingham Biomedical Services Unit. Ifng^mKate2^ mice were generated by Taconic and maintained at the University of Birmingham Biomedical Services Unit. Mice were culled between the ages of 8 and 18 weeks. Animals were used in accordance with Home Office Guidelines under a Project License awarded to D.R. Withers and approved by the University of Birmingham Animal Welfare and Ethical Review Body.

### Mouse tumor models

MC38 (kindly provided by Dr. Gregory Sonnenberg, Weill Cornell Medicine, New York, NY), MC38-Ova (obtained from AstraZeneca), CT26 (kindly provided by Professor Tim Elliot, University of Oxford, Oxford, UK), MC38^ptgs-/-^ (kindly provided by Dr Santiago Zelenay, University of Manchester, Manchester, UK) murine colon adenocarcinoma cells, and B16F10-Ova (obtained from AstraZeneca) murine melanoma cells were cultured in RPMI supplemented with 2mM L-glutamine (#21875034 Thermo Fisher Scientific), 10% FBS (#F9665 Sigma-Aldrich), and penicillin-streptomycin (#P4333 Sigma-Aldrich). EO771 (kindly provided by Dr. Fedor Berditchevski, University of Birmingham, Birmingham, UK), and 4T1 (obtained from AstraZeneca) mammary carcinoma cells were cultured in DMEM supplemented with 2mM L-glutamine (#21875034 Thermo Fisher Scientific), 10% FBS (#F9665 Sigma-Aldrich), and penicillin-streptomycin (#P4333 Sigma-Aldrich). All cells were cultured at 37°C with 5% CO_2_ before being harvested and suspended in Dulbeccos’ PBS (#D8662 Sigma-Aldrich) for tumor injection. 1 × 10^4^ 4T1, or 1 × 10^5^ EO771 tumor cells were injected in 50μl into the mammary fat pad of female mice under anesthesia via 2% gaseous isoflurane. 2.5 × 10^5^ (CT26, MC38, MC38-Ova, MC38^ptgs-/-^), or 5 × 10^5^ B16F10-Ova tumor cells were injected in 100μl subcutaneously into the pre- shaven left flank under anesthesia via 2% gaseous isoflurane. For orthotopic CT26 injections, 1 × 10^7^

CT26 tumor cells were injected in 25μl PBS into the colonic wall using an endoscope fitted with an integrated multi-purpose rigid telescope (Karl Storz Endoskope) under anesthesia via 2% gaseous isoflurane. Tumor size was periodically measured with a digital Vernier caliper, and the volume was calculated using the formula V = L x (W)^2^ x 0.52 in cubic millimeters, where L represents the longest diameter and W the perpendicular diameter to L for the tumor. Tumor weights were measured at the endpoint of the experiment. Mice were sacrificed 5h, 24h, 48h, or 72h post-photoconversion and tumors were harvested for analysis.

### Cell depletion

Depletion of NK cells and CD8^+^ T cells was achieved by administering anti-NK1.1 (PK136, 200μg) or anti-CD8α (53-6.7, 400μg), or InVivoMab mouse IgG2a isotype control (BioXCell, C1.18.4, 200μg), or InVivoMab rat IgG2a isotype control (BioXCell, 2A3, 400μg) in PBS via intraperitoneal injection every 3 days beginning on day 6 post tumor engraftment. Depletion of T_Regs_ was achieved by administering anti-OX40 (OX86, 200μg), or PBS via intraperitoneal injection on day 7 and 11 of tumor growth. Anti- NK1.1, anti-CD8α, and anti-OX40 antibodies were provided by AstraZeneca.

### Administration of IL-15:IL-15Rα complexes

rIL-15 (#210-15, Peprotech) and rIL-15Rα (#551-MR-100) were reconstituted in PBS at a ratio of 1:5 and incubated at 37°C for 30 minutes to form IL-15:IL-15Rα complexes. IL-15 complexes (2.5μg rIL- 15:12.5μg rIL-15Rα) were administered via intraperitoneal injections on day 7 and 12 of tumor growth.

### Splenic NK cell transfer

NK cells were enriched from the spleens of C57BL/6 Kaede mice using NK Cell Isolation Kit (#130-115- 818, Miltenyi Biotec) as per manufacturer’s instructions. Approximately 2.5x10^5^ splenic NK cells were transferred via intravenous injection into MC38 tumor bearing C57BL/6 mice 12 days after tumor engraftment, tissues were then collected 3 days later.

### Tumor compartment photo-labeling

Photoconversion of subcutaneous and mammary tumors were performed as previously described (*34*). Briefly, upon reaching 6-8mm in diameter both subcutaneous and mammary fat pad injected tumors were exposed to a 405-nm wavelength focused LED light (Dymax BlueWave QX4 fitted with 8mm focusing lens, DYM41572; Intertronics) using 9 cycles of 20 second light exposure with a 5- second break interval between each cycle, at a fixed distance. Black medium density fiberboard was used to shield the remainder of the mouse.

### Tissue dissociation

Tumors were processed as described previously Zhi et al. In short, tumors were cut into small pieces and enzymatically digested using 1mg/ml Collagenase D (#11088882001, Roche), and 0.1mg/ml DNase I (#101104159001, Roche) at 37°C for 20 - 22 minutes in a heat block shaker set to 1000rpm. Samples were then filtered through a 70-μm cell strainer, centrifuged for 5 mins at 400g and 4°C and resuspended in staining buffer (2%FBS and 2mM EDTA in PBS).

Spleens were crushed through a 70μm strainer, then incubated with Gey’s red blood cell lysis solution on ice for 5 minutes. Cells were harvested by centrifuging samples at 400g for 5 minutes at 4°C before resuspending cell pellets in staining buffer. Lymph nodes were collected, cleaned of fat, then cut into little pieces before incubating in 1mg/ml Collagenase D and 0.05mg/ml DNase I for 37°C in a heat block shaker at 1000rpm for 20 minutes. Samples were passed through a 70μm strainer, centrifuged at 400g and 4°C for 5 minutes before resuspending the cell pellet in staining buffer. Livers were cut into small 1-2mm pieces and pressed through a 100-μm, then washed through a 70μm strainer with RPMI. Liver suspensions were layered on top a 67% OptiPrep (#07820, STEMCELL Technologies) solution and centrifuged at 1000g for 25 minutes before collecting the interphase layer enriched for immune cells. Samples were washed again with RPMI, then centrifuged for 5 mins at 400g and 4°C before finally resuspending in staining buffer. Lungs were washed, cut into 1-2mm pieces before incubating tissue in 42.4μg/ml Liberase (#5401119001, Roche) and 0.02 mg/ml DNase I for 45 minutes at 37°C. Samples were passed through a 100-μm, then washed through a 70μm strainer with RPMI. Cells were pelleted by centrifuging at 400g and 4°C for 5 minutes, then cell pellets were resuspended in Gey’s solution and incubated on ice for 5 minutes. Samples were centrifuged for 5 minutes at 400g and 4°C before resuspending cell pellets in staining buffer. The small intestines were processed as described previously (*67*), briefly, fat and Peyer’s patches were removed for the small intestine, before cleaning internal contents and cutting tissue into small piece in Hank’s Balanced Salt Solution (HBSS) (#55037C, Sigma-Aldrich) containing 2% FBS. Next, the small intestines were shaken vigorously in HBSS with 2mM EDTA before incubating samples at 37°C for 20 minutes, then filtering through a nitex mesh, and washing thoroughly with HBSS. Samples were incubated with 1mg/ml collagenase VIII (#C2139, Sigma-Aldrich) and 0.1mg/ml DNase for 15 minutes, filtered through a 100μm, and then 70μm strainer before being centrifuged at 400g and 4°C for 5 minutes, and finally resuspended in staining buffer. Large intestines were processed similarly but digested in 0.85mg/ml Collagenase V (#C9263, Sigma-Aldrich), 1.25mg/ml Collagenase D, 1mg/ml Dispase (#17-105-041, Gibco), and 0.1mg/ml/ml DNase at 37°C for 45 minutes. Large intestines were then processed as per washing and straining steps used for small intestines, and finally cells were resuspended in staining buffer for use. All incubations were done in a shaking incubator set to 300rpm.

### In vitro assays

To assess NK cell mediated target cell killing, B16F0-Ova melanoma tumor cells were co-cultured with FACS NK cells as described previously (*68*). B16F10-Ova cells were incubated with 10μM Cell Trace Violet (#C34557, Invitrogen) for 15 minutes at 37°C, washed in RPMI and centrifuged at 350g for 5 minutes before plating 5x10^5^ cells/ml in complete RPMI supplemented with 10% FBS and 2mM L- glutamine. FACS sorted CD11b^+^ or CD49a^+^ NK cells from MC38 tumors were added to achieve 1:10, 1:20, and 1:40 Effector: Target ratios. Cells were co-cultured for 4 hrs at 37°C and 5% CO_2_.

To evaluate effect of PGE_2_ and TGF-β on NK cell phenotype and function, splenic NK cells were MACs enriched and cultured at 37°C and 5% CO_2_ in RPMI containing 300U rIL-2 (Peprotech), 50ng/ml rIL-15 (Peprotech) and either 0.1% DMSO (Sigma) or further supplemented with either 5ng/ml TGF-β, 43mg/ml 16,16-Dimethlyprostaglandin E2 (DiPGE_2_), or both TGF-β and DiPGE_2_ combined. NK cells were cultured for 48 hrs before restimulation with PMA and Ionomycin and subsequent flow analysis. In hypoxia experiments, MACs enriched NK cells were cultured with DMSO, TGF-β, or DiPGE_2_ as above or in a 1% O_2_ hypoxia chamber in parallel.

### Flow cytometry

To assess cytokine, granzyme and perforin production, cells were stimulated in 50 ng/ml Phorbol 12- myristate 13-acetate (PMA, #P1585, Sigma-Aldrich) and 1.5 µM ionomycin (#I0634, Sigma-Aldrich) for 4 hrs and in the presence of 10 µg/ml brefeldin A (#B6542, Sigma-Aldrich) added after the first hour. To assess cellular degranulation cells were stimulated like before, but 2μM Monensin (#420701, BioLegend) was used in place of Brefeldin A, and CD107a was added for the duration of the stimulation culture. Single cell suspensions underwent Fc blocking with anti-CD16/32 (2.4G2, BioLegend) in staining buffer on ice for 15 minutes before staining surface antigens in staining buffer on ice for 35 minutes. Cells were then fixed with BD CytoFix fixation buffer (#554655, BD Biosciences) on ice for 45 minutes before staining for intracellular markers overnight diluted in eBioscience permeabilization buffer (#00-8333-56, Thermo Fisher Scientific). Secondary intracellular staining to label Granzyme A and CCL5 was performed at room temperature for 45 minutes. Samples were transferred into staining buffer, with the addition of 1x10^4^ counting beads (#ACBP-100-10, Spherotech) being acquiring data on a BD LSR Fortessa X-20 (BD Bioscience) using FACSDiva 8.0.2 software (BD Bioscience) and analyzed with FlowJo v10 (BD Bioscience). Surface and intracellular antibodies used were against the following mouse antigens: CCL5 purified (Goat IgG, R&D Systems), CD107a BV786 (1D4B, BioLegend), CD11b PE-Dazzle594 (M1/70, BioLegend), CD200r1 APC (OX-110, BioLegend), CD3 BV711 or BUV737 (17A2, BioLegend), CD45 FITC or BV510 (30-F11, BioLegend), CD49a BUV395 (RM4-5, BD Bioscience), CD49b BV711(DX5, BioLegend), CD69 PE-Cy7 or BV711 (H1.2F3, BioLegend), DNAM-1 BV605 (TX42.1, BioLegend), EOMES PE or PE-Cy7 (Dan11mag, BioLegend), Granzyme A purified (3G8.5, BD Biosciences), Granzyme B BV421 (QA18A28, BioLegend), Granzyme C PE-Cy7 (SFC108, BioLegend), IFNγ BUV737 (XMG1.2, BioLegend), IgG2b BV605 or R718 (R12-3, BioLegend), IL-7Rα BV605 or BV421 (AFR34, BioLegend), Ki-67 BV711 or AF700 (SolA15, BioLegend), KLRG1 BV605 (2F1, BioLegend), LAG3 BV786 (C9B7W, BioLegend), NK1.1 BV650 or BV786 (PK136, BioLegend), NKG2A/C/E BV421 (20d5, BioLegend), NKp46 BV650 or BV786 (29A1.4, BioLegend), OX40 BV711 (OX-86, BioLegend), PD-1 PE-Cy7 (RMP1-30, BioLegend), Perforin APC (S16009A, BioLegend), T-bet e660 (eBio4B10, BioLegend), and TCRβ BUV737 (H57-597, BioLegend). AF647-conjugated polyclonal donkey anti-goat IgG (#A32849, Thermo Fisher Scientific) was used to identify and amplify CCL5 staining.

### FACS isolation

Single cell suspensions were stained for antibodies raised against the following antigens: CD45 FITC (30-F11, BioLegend), CD11b PE-Dazzle594 (M1/70, BioLegend), TCRβ BUV737 (H57-597, BioLegend), NKp46 BV650 (29A1.4, BioLegend), and CD49a BUV395 (RM4-5, BD Biosciences), in addition to Live/Dead Near-IR viability stain (#L10119, Thermo Fisher Scientific). NK cells (Live CD45^+^ CD11b^-/low^, TCRβ^-^ CD3^-^ NKp46^+^) were sorted into CD11b^+^ CD49a^-^ and CD11b^-^ CD49a^+^ cells using a FACS Aria II Cell sorter (BD Biosciences) and then used for *in vitro* assays. For scRNA-seq, singles cells were stained using antibodies against the following: CD45-BUV395 (30-F11, BioLegend), CD11b-APC (M1/70, BioLegend), Ter119-PE-Cy7 (TER-119, BioLegend), NK1.1-BV650 (PK136, BioLegend). Tumor infiltrating lymphocytes (Live CD45^+^ Ter119^-^ CD11b^-/low^) were sorted into two groups based on the presence of the Kaede Red, namely, G48 and R48 referring to Kaede Green 48h and Kaede Red 48h.

### Single-cell library construction and sequencing

Gene expression libraries from were prepared from FACS-sorted populations of single cells using the Chromium Controller and Chromium Single Cell 3’ GEM Reagent Kits v3 (10x genomics, Inc.) according to the manufacturer’s protocol. The resulting sequencing libraries comprised of standard Illumina paired-end constructs flanked with P5 and P7 sequences. The 16 bp 10x barcode and 10 bp UMI were encoded in read 1, while read 2 was used to sequence the cDNA fragment. Sample index sequences were incorporated as the i7 index read. Paired-end sequencing (2 x 150 bp) was performed on the Illumina NovaSeq 6000 platform. The resulting .bcl sequence data were processed for QC purposes using bcl2fastq software (v2.20.0.422) and the resulting fastq files were assessed using FastQC (v0.11.3), FastqScreen (v0.9.2) and FastqStrand (v0.0.5) prior to alignment and processing with the CellRanger (v6.1.2) pipeline.

### Processing of scRNA-seq

Single-cell gene expression data from CellRanger count output (filtered features, barcodes, and matrices) were analyzed using the Scanpy (*69*) (v1.8.2) workflow. Doublet detection was performed using Scrublet (*70*) (v0.2.1), with cells from iterative sub-clustering flagged with outlier Scrublet scores labelled as potential doublets. Cells with counts mapped to >6000 or <1000 genes were filtered. The percentage mitochondrial content cut-off was set at <7.5%. Genes detected in fewer than 3 cells were filtered. Total gene counts for each cell were normalized to a target sum of 10^4^ and log1p transformed. There resulted in a working dataset of 46,342 cells. Next, highly variable features were selected based on a minimum and maximum mean expression of ≥0.0125 and ≤3 respectively, with a minimum dispersion of 0.5. Total feature counts, mitochondrial percentage, and cell cycle scores, where indicated, were regressed out. The number of principal components used for neighborhood graph construction was set to 50 initially, and subsequently 30 for subgroup processing. Clustering was performed using the Leiden algorithm with resolution set between 0.8 and 1.0. Uniform manifold approximation and projection (UMAP, v0.5.1) was used for dimensional reduction and visualization, with a minimum distance of 0.3, and all other parameters according to the default settings in Scanpy.

### Analysis of scRNA-seq from mouse tumor models

Cell types of interest were subset and re-clustered as described above. Resulting clusters were annotated using canonical marker genes. Gene set scoring was performed using Scanpy’s tl.score genes tool. Gene sets were obtained from the Molecular Signature Database (MSigDB) inventory, specifically KEGG or Gene Ontology (GO), using the R package msigdbr (v7.5.1) or published RNAseq data. Differential gene testing was performed using the Wilcoxon rank sum test implemented in Scanpy’s tl.rank_genes_groups. Trajectory analysis was performed using partition-based graph abstraction (PAGA) (*71*) and diffusion pseudotime (*72*) implemented in Scanpy. For pseudotime analysis, the differentiation trajectory was rooted in the Kaede-green dominant cluster, which represents the cellular subset most associated with newly entering cells. Calculation of differential expression across pseudotime was performed using the tradeSeq framework (*38*). The 24h and 72h scRNA-seq data was processed as previously described (*34*). Integration and label transfer of 24h and 72h scRNA-seq data and the main data set was performed using Scanpy’s tl.ingest tool.

### Analysis of RNA-seq from human tumors

The cancer genome atlas (TCGA) bulk-RNAseq data of human colorectal adenocarcinoma (TCGA-COAD) was accessed using TCGAbiolinks (*45*). Cellular deconvolution was performed using Cibersortx (*73*). scRNA-seq from human colorectal tumors (*46*) was analyzed using the Scanpy (v1.8.2) workflow and analyses as outlined above. Filtering for quality control was performed according to the parameters outlined in the original publications. Batch integration was performed using the Harmony algorithm. Cell annotations from original publications were checked and refined using canonical marker gene expression. Gaussian kernel density estimation to compute density of cells from various conditions in the UMAP embedding was performed using Scanpy’s tl.embedding_density. Pre-ranked gene set enrichment analysis (GSEA) was implemented in fgsea (v1.24), using the Wald statistic as the gene rank metric.

### Statistical analysis

Data collected were analyzed using Flow Jo 10.8.1 software (BD BioScience) and GraphPad Prism 9.4.0. UMAPs generated in FlowJo were made using DownSampleV3 and UMAP plugins, both available on the FlowJo Exchange. Normality was determined using the Shapiro-Wilk test. Pairs of samples were compared using an unpaired two-tailed Mann-Whitney T test. When comparing more than two sets of data, statistical significance was determined by either one-way ANOVA with Tukey’s multiple comparison test, or two-way ANOVA with Šidák’s multiple comparisons test. Two data points were identified as outliers by the ROUT method in Supplemental figure 11E-F and removed from analysis. All graphs show mean ± SD unless stated otherwise. *p≤0.05, **p≤0.01, ***p≤0.001.

## Acknowledgements

We thank Dr Y. Miwa (Tsukuba University) and Dr O. Kanagawa (RCAI, RIKEN) and Dr. M. Tomura (Osaka Ohtani University) for the Kaede mice. We thank Dr S. Zelenay for kindly sharing MC38^ptgs-/-^ cells. We thank the Biomedical Services Unit at the University of Birmingham for all their help with *in vivo* experimental work. We thank A. Ptasinska and colleagues at Genomics Birmingham, the genomic and sequencing facility of the University of Birmingham, for their help with single cell RNA sequencing and the University of Birmingham Flow Cytometry Platform. We thank Gareth Howell and the University of Manchester flow cytometry core, and Andy Hayes and Claire Morrisroe in the University of Manchester Genomic Technologies core facility for their help with single cell RNA sequencing. Results shown here are in part based upon data generated by the TCGA Research Network: https://www.cancer.gov/tcga.

## Data and Materials Availability

The scRNA-seq data have been made available on the GEO public repository under accession numbers GSE221064. Published scRNA-seq data and accompanying metadata was accessed and downloaded from the GEO repository using the ascension number GSE178341. All (other) data needed to evaluate the conclusions in the paper are present in the paper or the Supplementary Materials.

## List of Supplementary Material

Fig. S1. Single cell RNA-sequencing of Tumor Infiltrating Lymphocytes 48 hrs after photoconversion Fig. S2. Tracking transcriptomic changes in tumor infiltrating NK cells over time

Fig. S3. The CD3- NK1.1+ compartment of tumors is phenotypically distinct to that of other tissues Fig. S4. Changes in NK cell integrin expression over time observed in multiple tumor models

Fig. S5. Limited NK cell egress from tumors

Fig. S6. Confirmation of changes in NK cell function over time in the TME across multiple pre-clinical tumor models

Fig. S7. Transferred splenic NK cells upregulate CD49a and Granzyme C expression after entry into MC38 tumors but not healthy tissues

Fig. S8. Investigating the contributions of TGF-β and PGE_2_ *in vitro*, and Tregs *in vivo, in* suppressing NK cell effector functions

Fig. S9. Further analysis of human CRC data

Fig. S10. Gating strategies for identification of human NK cells

Fig. S11. Depletion of NK cells after subcutaneous tumors have become established has minimal impact on tumor growth

Fig. S12. Administration of IL-15:IL-15Rα complexes enhance formation of CD11b+ CD49a+ intratumoral NK cells across multiple tumor models

