## Supplemental Data for "Rapid establishment of a tumor-retained state curtails the contribution of conventional NK cells to anti-tumor immunity in solid cancers"

**Supplementary Materials**

**Fig. S1. Single cell RNA-sequencing of Tumor Infiltrating Lymphocytes 48 hrs after photoconversion.** (A) Cartoon summarizing experimental plan to isolate photo-labelled TILs from MC38 tumors for analysis by scRNA-seq. (B) Gating strategy used to isolate TILs and ensure appropriate capture of NK cells. (C) UMAP showing immune cell clusters within the total TIL compartment. (D) UMAP showing the distribution of Kaede Green+ and Kaede Red+ cells amongst the immune cell clusters (E) Dot plots showing canonical marker expression used to initially define major cell clusters. (F) Dot plots showing differentially expressed genes defining the 8 clusters comprising the initial main 'NK cell' cluster in 'C'. (G) UMAPs summarizing expression of key genes used to define the NKT cluster. (H) UMAPS summarizing expression of key genes used to define the ILC cluster. (I) Violin plots showing expression of *Tbx21*, *Eomes*, and *Rora* across the clusters.

**Fig. S2. Tracking transcriptomic changes in tumor infiltrating NK cells over time**

To further investigate how NK cells change over time within the TME, scRNA-seq data generated from FACS-isolated TILs at two time points post photoconversion of MC38 tumors (*34*) was reanalyzed (data set E-MTAB-10176). This data set contained 4 samples, comprised of Kaede Green+ and Kaede Red+ TILs each at 24 and 72 hrs post photoconversion. (A) UMAPs showing the three NK clusters identified in the data set (NK_a, NK_b, NK_c) and the pseudotime trajectory rooted in NK_a. (B) Proportion of NK cells from the different samples in each cluster, with NK_a almost entirely dominated by Kaede Green+ cells and NK_c comprised of only Kaede Red+ cells. (C) UMAPs showing the distribution of cells in each sample across the three clusters. (D) Dot plots showing expression of selected genes used to further characterize the clusters. Note, this is the same genes assessed in Fig. 1D. (E) UMAPs showing expression of selected genes across the NK clusters, highlighting differential expression of integrins, granzymes and *Ccl5*.

**Fig. S3. The CD3- NK1.1+ compartment of tumors is phenotypically distinct to that of other tissues.** To compare the phenotype of NK cells within tumors to the NK and ILC1 cells found in healthy tissue, flow cytometry was used to assess cells isolated from MC38 and B16-F10 tumors as well as spleen, liver, small intestine, and colon. (A) Table showing the ILC and NK cell 'markers' analyzed. (B) Flow cytometry plot showing gating on CD3- NK1.1+ cells (amongst live CD45+ cells). (C) UMAP showing 4,500 $CD3^-$ $Nk1.1^+$ cells from each tissue in non-tumor bearing mice (n=3), and from MC38 and B16F10 tumors (n=3). (D) UMAPS showing the distribution of cells in each tissue. UMAPs showing expression of proteins typically associated with ILCs rather than NK cells (IL-7R$\alpha$, DNAM-1, CXCR6, CD200r1, CD49a). (F) UMAPs showing expression of proteins typically associated with NK cells (T-bet, Eomes, CD49b, CD11b). (G) UMAP summarizing the distribution of NK cells versus non-NK ILCs. (H) The proportion of cells within the CD3- NK1.1+ gate that are NK cells within each tissue, defined in 'G'. (I) Graph showing MFI of selected proteins in the non-NK ILC and NK regions of the UMAP defined in 'G'. Statistical significance was determined by two-way ANOVA with Šidák's multiple comparisons test. *P<0.05, ****P<0.0001.

**Fig. S4. Changes in NK cell integrin expression over time observed in multiple tumor models.** The changes in NK cell integrin expression over time in the TME observed in MC38 tumors were further assessed in CT26 and E0771 tumors, grafted subcutaneously into BALB/c Kaede and into the mammary fat pad of C57BL/6 Kaede mice respectively. (A) Expression of CD49a versus CD11b by Kaede Green+ and Kaede Red+ NK cells (CD3- NKp46+) isolated from CT26 tumors at 5 (n=5), 24 (n=5), and 72 (n=6) hrs post photoconversion. Data shown for 1 independent repeat at 5 hrs and 24 hrs, and 2 pooled independent repeats at 72 hrs post-labeling. (B) The proportion of Kaede Green/Red for each NK cell subset at each time point post photoconversion. (C) Total number of Kaede Green+ and Kaede Red+ NK cells at each time point post photoconversion. (D) Expression of CD49a versus CD11b by Kaede Green+ and Kaede Red+ NK cells (CD3- NK1.1+) isolated from E0771 tumors at 24 (n=9) and 72 (n=4) hrs post photoconversion. Data shown for 2 pooled independent repeats at 24 hrs, and 1 independent repeat at 72 hrs post-labeling. (E) The proportion of Kaede Green/Red for each NK cell subset at each time point post photoconversion. (F) Total number of Kaede Green+ and Kaede Red+ NK cells at each time point post photoconversion.

**Fig. S5. Limited NK cell egress from tumors.**

To investigate the egress of NK cells from tumors, MC38 and CT26 tumors were photoconverted and the Kaede Red+ cells within the draining (inguinal) lymph node and spleen assessed at 24 and 72 hrs post photoconversion. (A) Identification of Kaede Red+ NK cells (CD3- NK1.1+) in the spleen and dLN of mice bearing MC38 tumors, with CD49a versus CD11b expression used to assess NK cell phenotype. (B) Total number on NK cells within the spleen and dLN at 24 (n=7) and 72 (n=8) hrs post photoconversion, showing NK cell subsets defined by CD49a versus CD11b expression. Data pooled from 2 independent experiments. (C) Proportion of NK cells within the spleen and dLN at 24 and 72 hrs post photoconversion in each NK cell subset defined by CD49a versus CD11b expression. (D) Identification of Kaede Red+ NK cells (CD3- NKp46+) in the spleen and dLN of mice bearing CT26 tumors, with CD49a versus CD11b expression used to assess NK cell phenotype. (E) Total number on NK cells within the spleen and dLN at 24 (n=4) and 72 (n=7) hrs post photoconversion, showing NK cell subsets defined by CD49a versus CD11b expression. (F) Proportion of NK cells within the spleen and dLN at 24 and 72 hrs post photoconversion in each NK cell subset defined by CD49a versus CD11b expression. CT26 Data from 1 independent experiment. Statistical significance was determined by two-way ANOVA with Šidák's multiple comparisons test. *P<0.05, **P<0.01, ***P<0.001, ****P<0.0001.

**Fig. S6. Confirmation of changes in NK cell function over time in the TME across multiple pre-clinical tumor models.** (A) Representative expression of mKate2 as a read-out for *Ifng* expression by NK cells versus T cells after *ex vivo* culture of splenocytes ± IL-12 and IL-18 (n=2). (B) Cartoon showing experimental set up with CT26 tumors grafted subcutaneously on the flank of BALB/c Kaede mice and photoconverted 12 days later. (C) Proportion of NK cells producing CCL5, CD107a, Granzyme A, Granzyme B, Granzyme C, and Perforin after *ex vivo* stimulation at 24 hrs post photoconversion. Data representative of 2 independent experiments, CCL5 n=4, all other parameters n=5. (D) Cartoon showing experimental set up with 4T1 tumors grafted into the mammary fat pad (m.f.p) of BALB/c Kaede mice and photoconverted 12 days later. (E) Proportion of NK cells producing CCL5, Granzyme A, Granzyme B, Granzyme C, and Perforin after *ex vivo* stimulation at 24 hrs post photoconversion. Data was pooled from 2 independent experiments, n=6 at each time point. (F) Cartoon showing experimental set up with CT26 tumors grafted into the colon wall of C57BL/6 Kaede mice via endoscopy. (G) Endoscopic photo of the CT26 tumor growing within the colon 12 days after injection. (H) Flow cytometry plots showing identification of CXCR6+ ILC and Eomes+ NK cells within the CD3- NKp46+ population within the tumor. Further analysis of CD49a versus CD11b revealed the presence of three subsets within the Eomes+ NK cells, consistent with analysis of CT26 tumors grown subcutaneously. (I) MFI of Granzyme A, Granzyme C and Perforin expression. Data from a single independent experiment, n=4. Statistical significance was determined by two-way ANOVA with Šidák's multiple comparisons test (C and E), and one-way ANOVA with Tukey's multiple comparison test comparing all means (I). *P<0.05, **P<0.01, ***P<0.001, ****P<0.0001.

**Fig. S7. Transferred splenic NK cells upregulate CD49a and Granzyme C expression after entry into MC38 tumors but not healthy tissues.** Approximately 250,000 MACS-enriched NK cells isolated from the spleens of C57BL/6 Kaede mice and injected intravenously into wild-type C57BL/6 mice bearing MC38 tumors (n=4). Three days after transfer, mice were culled and cells isolated from spleen, liver, lung, and MC38 tumors analyzed after *ex vivo* restimulation. (A) Cartoon summarizing the experimental design. (B) Flow cytometry plots showing efficiency of NK cell MACS-enrichment. (C) Flow cytometry plots showing identification of transferred Kaede Green+ NK cells in different tissues. Proportion of transferred NK cells expressing CD11b (D), CD49a (E),g ranzyme A (F) and Granzyme C (G) in each tissue. Data representative of 2 independent experiments. Statistical significance was determined by one-way ANOVA with Dunnett's multiple comparison test comparing means to MC38. *P<0.05, **P<0.01, ***P<0.001, ****P<0.0001.

**Fig. S8. Investigating the contributions of TGF-β and $PGE_2$ *in vitro*, and Tregs *in vivo, in* suppressing NK cell effector functions.** TGFβ, PGE2, and hypoxia were investigated *in vitro* and *in vivo* as potential mechanisms promoting NK cell differentiation to the CD49a expression tumor retained state. Splenic NK cells isolated and then cultured with IL-2/IL-15 further supplemented with TGF-β and/or $PGE_2$ for 48 hrs, restimulated *ex vivo* and analyzed by flow cytometry. (A) Bar chart showing proportion of CD11b+ and CD49a+ NK cells after 48 hrs in culture in RPMI with IL-2/IL-15 further supplemented with TGF-β and/or DiPGE2. (B) Bar charts showing proportion of CCL5, IFNγ, NKG2A, Granzyme A, and Granzyme C expressing NK cells. Splenic NK cells were isolated and cultured with +IL-2/IL-15, in addition to TGF-b or DiPGE2 in either a 1% O2 hypoxic chamber or normal incubator. After 2 days cells were then restimulated with PMA and Ionomycin and analyzed. (C) Bar chart comparing proportion of CD11b+ NK cells, and CD49a+ cells. Representative histograms alongside proportion of NK cells positive for (D) NKG2A, (E) CCL5, (F) Granzyme A, or (G) Granzyme C. (H) Cartoon summarizing experimental design for depleting Tregs in MC38 tumors to impede TGFβ production *in vivo*. WT C57BL/6 mice treated with PBS (n=11) or anti-OX40 antibodies (OX-86, mIgG2A, n=7) on D7 and D11, then analyzed 2 days later. (I) Tumor weight. (J) Enumeration of different TIL populations, showing CD4+ FoxP3- T cells, FOXP3+ Tregs and NK cells. (K) Proportion of NK cells in CD11b+ CD49a-, CD11b- CD49a- and CD11b- CD49a+ subsets. (L) Proportion of NK cells producing CCL5, IFNγ, Granzyme A, Granzyme C after *ex vivo* restimulation. Statistical significance was determined by one-way ANNOVA with Tukey's multiple comparison test (A and B), two-way ANNOVA with Šidák's multiple comparison test (C-G, and L), or unpaired t-tests (I and J). *P<0.05, **P<0.01, ***P<0.001, ****P<0.0001.

**Fig. S9. Further analysis of human CRC data.** (A) CIBERSORTx deconvolution of immune cells from bulk transcriptomes of 521 human colorectal cancers, from TCGA. (B) Further subdivision of NK cells from deconvolution results in (A). (C) UMAP of 71,318 T, NK or ILC cells from scRNA-seq of 62 human CRC samples, from GSE178341, and (D) canonical marker gene expression. (E) Gene set enrichment for GO:BP Natural killer cell activation between NK1 clusters. (F) Gene set enrichment analysis (GSEA) of KEGG Natural killer cell-mediated cytotoxicity between scRNA-seq of NK cells from tumor and normal adjacent tissue, in human CRC.

**Fig. S10. Gating strategies for identification of human NK cells.**

Representative plots showing gating strategy for NK cell subsets from (A) primary CRC tumor, (B) unaffected colon tissue, and (C) blood.

**Fig. S11. Depletion of NK cells after subcutaneous tumors have become established has minimal impact on tumor growth.** To assess the contribution of NK cells to the control of established tumors, MC38 or B16-F10 tumors were engrafted subcutaneously on the flank of mice and administered with anti-NK1.1 depleting or isotype control antibodies from day 6 of tumor growth. (A) Tumor growth curves of MC38 tumors treated with anti-NK1.1 or IgG control antibodies (indicated by red arrows). (B) Tumor weight upon tissue harvest. (C) Flow cytometry plots showing efficient depletion of NKp46+ CD49b+ cells in the MC38 tumors with anti-NK1.1 antibodies. (D) Enumeration of the number of NK cells present within MC38 tumors. Data from one independent experiment, n=5. (E) Flow cytometry histogram showing MHCI expression on MC38 and B16-F10 cell lines +/- IFN$\gamma$, alongside MFI, n=3. (F) Tumor growth curves of B16-F10 tumors grafted into WT or Rag-/- mice and treated with anti-NK1.1 or IgG control antibodies (indicated by red arrows), n=3 for Rag-/- mice treated with anti-NK1.1. and n=4 for all other conditions. (G) Tumor weight upon tissue harvest. (H) Flow cytometry plots showing efficient depletion of NKp46+ CD49b+ cells in the MC38 tumors with anti-NK1.1 antibodies. (I) Flow cytometry plots depicting identification of CD11c+ MHCII+ DC within B16F10 tumors from WT and RAG2 mice. (J) Enumeration of the number of NK cells present within MC38 tumors. (K) Enumeration of intratumoral DC normalized to B16F10 tumor weight in each treatment group. Statistical significance was determined by unpaired *t*-tests (B and D), two-way ANOVA with Šidák's multiple comparisons test (E), and one-way ANOVA with Tukey's multiple comparison test (F and H) comparing means of pre-selected pairs (IgG control to anti-NK1.1, and RAG IgG to RAG anti-NK1.1). *P<0.05, **P<0.01, ***P<0.001, ****P<0.0001.

**Fig. S12. Administration of IL-15:IL-15Rα complexes enhances tumor control in MC38 and B16-F10 models.** MC38 tumor cells were engrafted into C57BL/6 Kaede mice and administered with either PBS or IL-15:IL-15Rα complexes on day 7 and 12 of tumor growth. (A) Cartoon diagram showing experimental setup. (B) MC38 tumor growth curve from 1 independent experiment (PBS n=5, IL-15:IL-15Rα n=4). (C) Representative flow cytometry plots showing CD11b and CD49a integrin expression on NK cells (CD3- NK1.1+) isolated from MC38 tumors. (D) Proportion of NK cells with CD11b+CD49a-, CD11b-CD49a-, CD11b-CD49a+, and CD11b+CD49a+ phenotype. (E) Representative flow cytometry plot showing Kaede Green versus Kaede Red expression by NK cells, alongside enumeration of % Kaede Green+ NK cells in either PBS (n=4) or IL-15:IL-15Rα (n=3) treatment. (F) Proportion of Kaede Red+ NK cells across NK cell subsets in either treatment. (G) Representative histograms and enumeration of Granzyme A, Granzyme B, Granzyme C, Perforin, and CCL5 production (MFI) by NK cells from MC38 tumors after *ex vivo* stimulation. (H) Cartoon showing experimental setup whereby B16-F10 tumors were engrafted into WT C57BL/6 mice and administered with i.p. injection of PBS or IL-15:IL-15Rα complexes on day 7 and 12 of tumor growth. (I) B16-F10 tumor growth from 1 independent experiment (PBS n=5, IL-15:IL-15Rα n=5). (J) Representative flow cytometry plots displaying CD11b versus CD49a expression, and (K) enumeration of each NK cell subset comprising intratumoral NK cells. (L) Enumeration of CCL5, Granzyme A, Granzyme B, Granzyme C, and Perforin production (MFI) by NK cells from B16F10 tumors after *ex vivo* stimulation. Statistical significance was determined by two-way ANOVA with Šidák's multiple comparisons test (B, D, F, G, I, K, L, and M), and unpaired t-test (E). *P<0.05, **P<0.01, ***P<0.001, ****P<0.0001.

**Fig S1.**

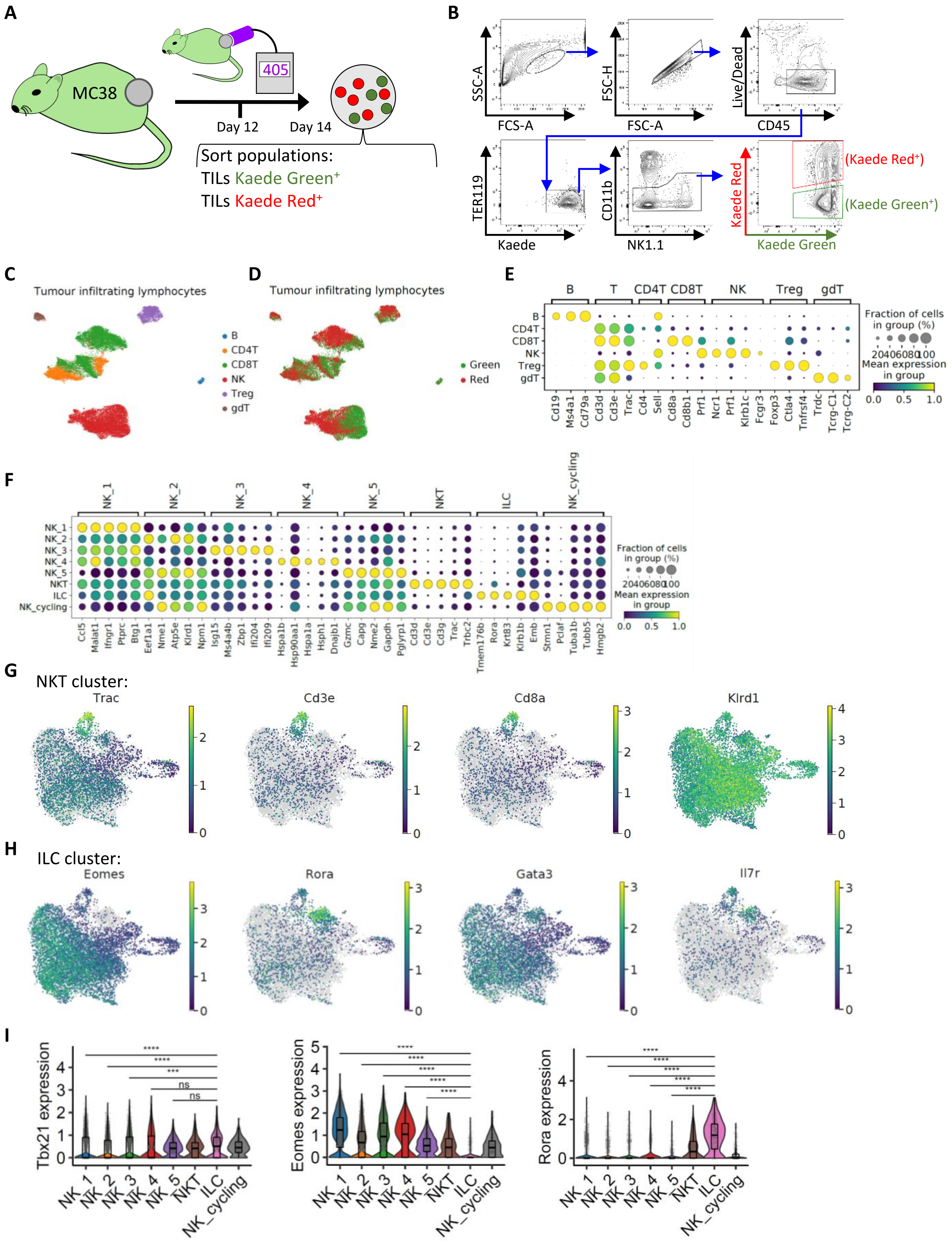
A
MC38
405
Day 12
Day 14
Sort populations:
TILs Kaede Green+
TILs Kaede Red+
B
SSC-A
FCS-A
FSC-H
FSC-A
Live/Dead
CD45
TER119
Kaede
CD11b
NK1.1
Kaede Red
Kaede Green
(Kaede Red+)
(Kaede Green+)
C
Tumour infiltrating lymphocytes
B
CD4T
CD8T
NK
Treg
gdT
D
Tumour infiltrating lymphocytes
Green
Red
E
B
T
CD4T
CD8T
NK
Treg
gdT
Cd19
Ms4a1
Cd79a
Cd3d
Cd3e
Trac
Cd4
Sell
Cd8a
Cd8b1
Prf1
Ncr1
Prf1
Klrb1c
Fcgr3
Foxp3
Ctla4
Tnfrsf4
Trdc
Tcrg-C1
Tcrg-C2
Fraction of cells in group (%)
20 40 60 80 100
Mean expression in group
0.0
0.5
1.0
F
NK_1
NK_2
NK_3
NK_4
NK_5
NKT
ILC
NK_cycling
Ccl5
Malat1
Ifngr1
Ptprc
Btg1
Eef1a1
Nme1
Atp5e
Klrd1
Npm1
Isg15
Ms4a4b
Zbp1
Ifi204
Ifi209
Hspa1b
Hsp90aa1
Hspa1a
Hsph1
Dnajb1
Gzmc
Capg
Nme2
Gapdh
Pglyrp1
Cd3d
Cd3e
Cd3g
Trac
Trbc2
Tmem176b
Rora
Krt83
Klrb1
Emb
Stmn1
Pclaf
Tuba1b
Tubb5
Hmgb2
Fraction of cells in group (%)
20 40 60 80 100
Mean expression in group
0.0
0.5
1.0
G
NKT cluster:
Trac
Cd3e
Cd8a
Klrd1
H
ILC cluster:
Eomes
Rora
Gata3
Il7r
I
Tbx21 expression
Eomes expression
Rora expression
****
***
ns
NK_1
NK_2
NK_3
NK_4
NK_5
NKT
ILC
NK_cycling

**Fig S2.**

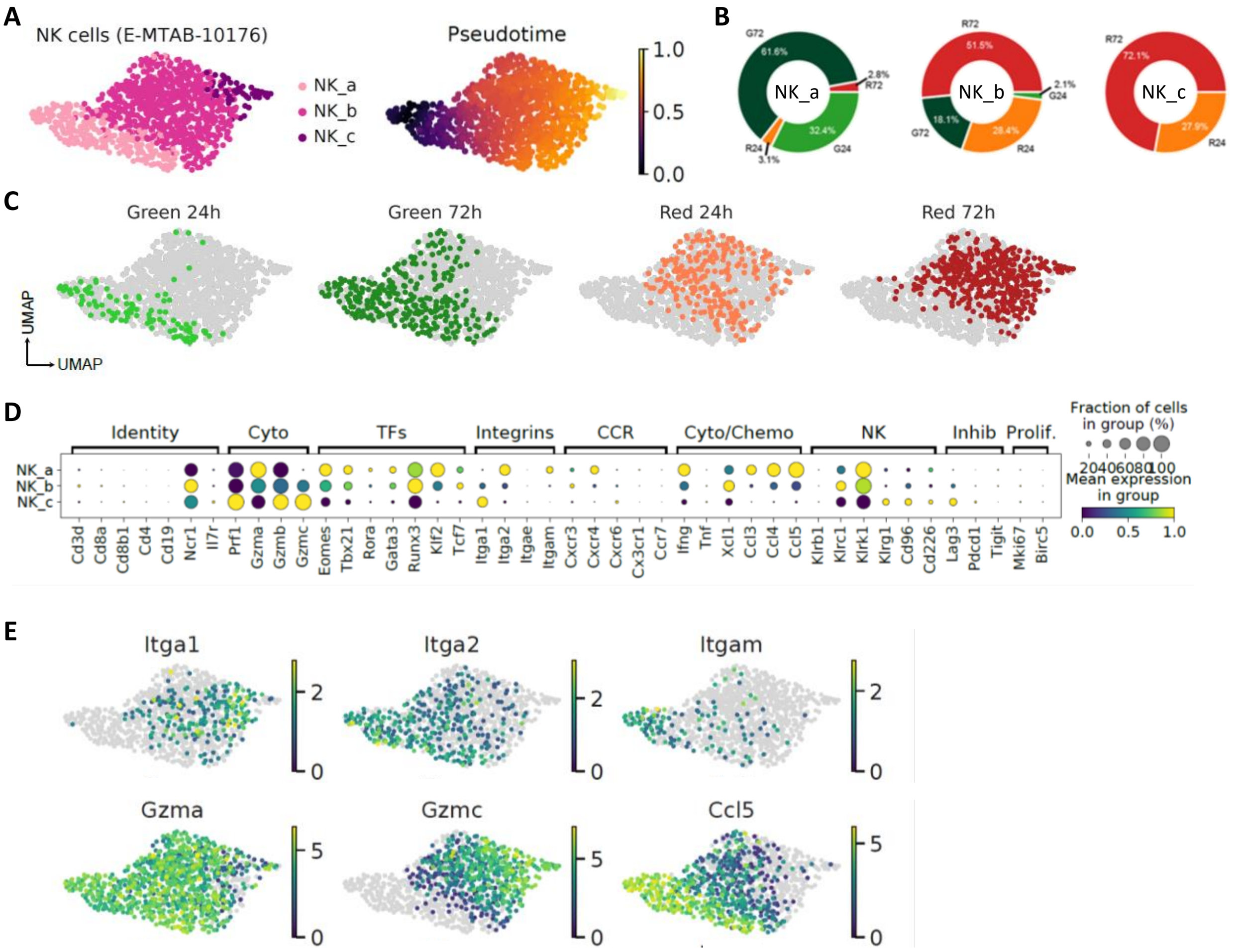
A
NK cells (E-MTAB-10176)
NK_a
NK_b
NK_c
Pseudotime
1.0
0.5
0.0
B
NK_a
NK_b
NK_c
C
Green 24h
Green 72h
Red 24h
Red 72h
UMAP
UMAP
D
Identity
Cyto
TFs
Integrins
CCR
Cyto/Chemo
NK
Inhib
Prolif.
Fraction of cells in group (%)
Mean expression in group
E
Itga1
Itga2
Itgam
Gzma
Gzmc
Ccl5

**Fig S3.**

**A**

| Non-NK ILC markers | NK Markers |
|---|---|
| T-bet | T-bet |
| IL-17Rα | Eomes |
| DNAM-1 | CD49b |
| CXCR6 | CD11b |
| CD200r1 | |
| CD49a | |

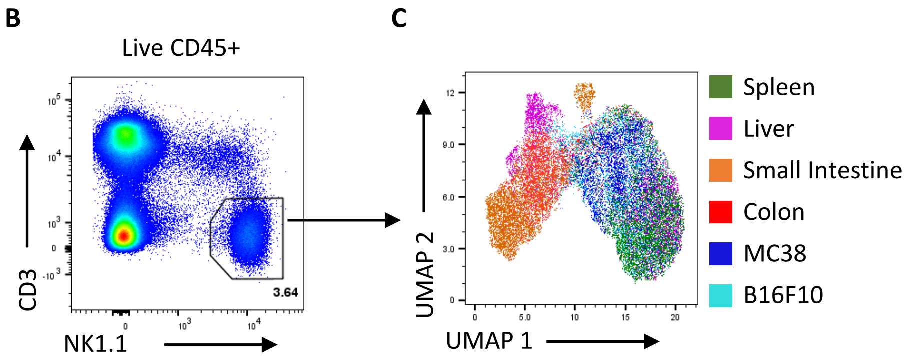


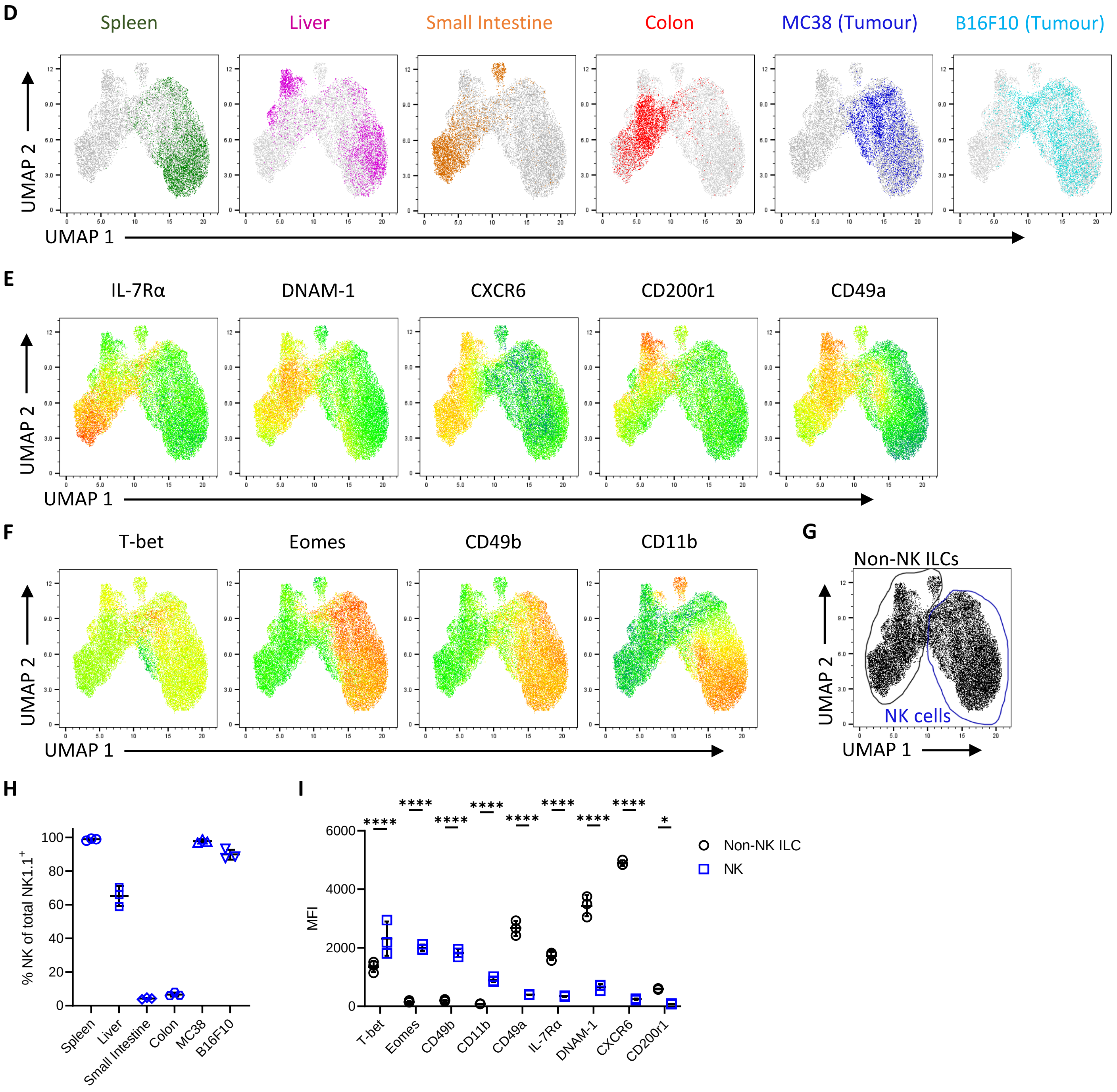

Fig S4.

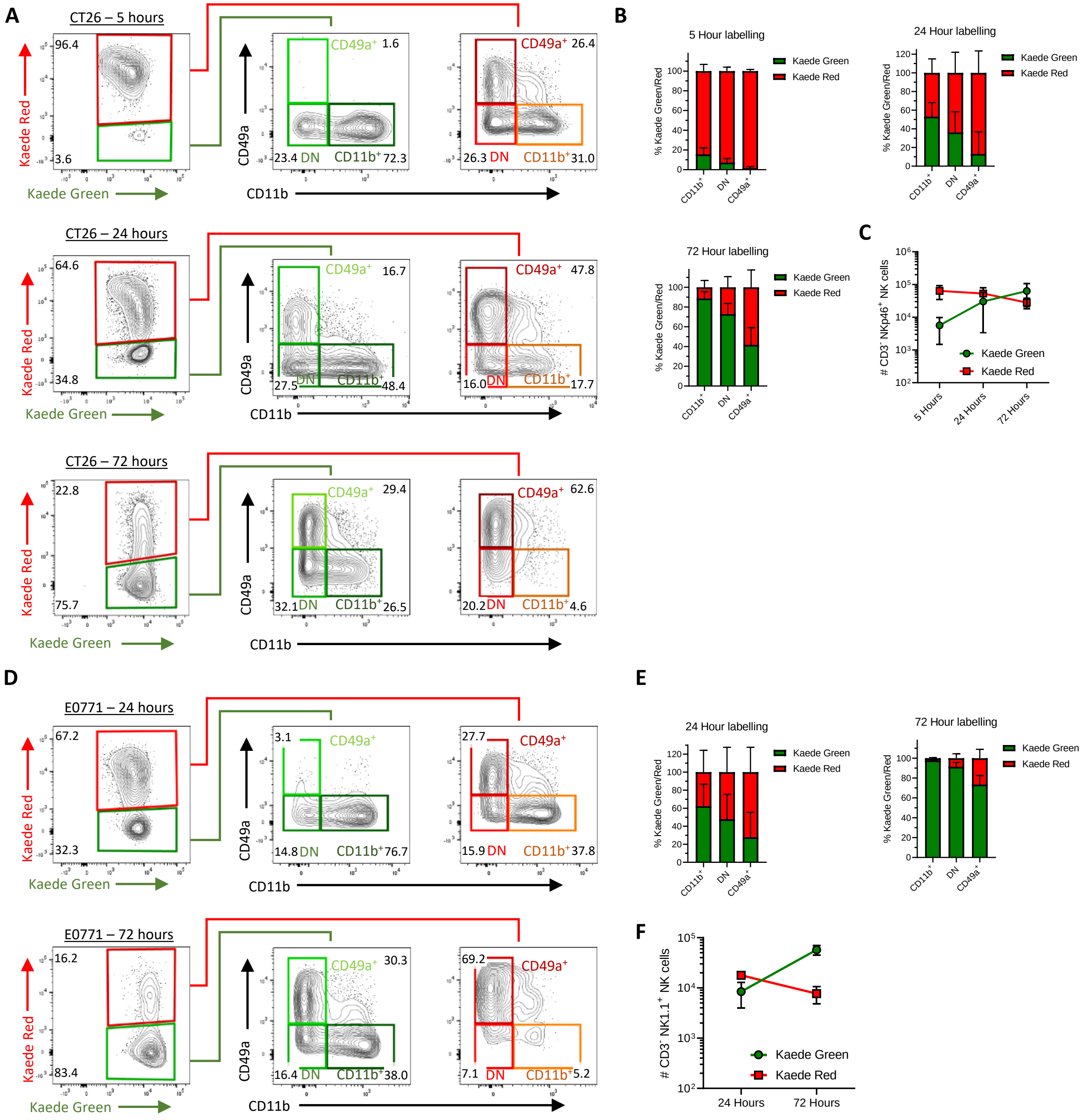
A
CT26 – 5 hours
Kaede Red
Kaede Green
96.4
3.6
CD49a
CD11b
CD49a+ 1.6
23.4 DN
CD11b+ 72.3
CD49a+ 26.4
26.3 DN
CD11b+ 31.0
CT26 – 24 hours
64.6
34.8
CD49a+ 16.7
27.5 DN
CD11b+ 48.4
CD49a+ 47.8
16.0 DN
CD11b+ 17.7
CT26 – 72 hours
22.8
75.7
CD49a+ 29.4
32.1 DN
CD11b+ 26.5
CD49a+ 62.6
20.2 DN
CD11b+ 4.6
B
5 Hour labelling
24 Hour labelling
72 Hour labelling
% Kaede Green/Red
Kaede Green
Kaede Red
CD11b+
DN
CD49a+
C
### CD3- NKp46+ NK cells
5 Hours
24 Hours
72 Hours
D
E0771 – 24 hours
67.2
32.3
3.1
CD49a+
14.8 DN
CD11b+ 76.7
27.7
CD49a+
15.9 DN
CD11b+ 37.8
E0771 – 72 hours
16.2
83.4
CD49a+ 30.3
16.4 DN
CD11b+ 38.0
69.2
CD49a+
7.1 DN
CD11b+ 5.2
E
24 Hour labelling
72 Hour labelling
F
### CD3- NK1.1+ NK cells
24 Hours
72 Hours

**Fig S5.**

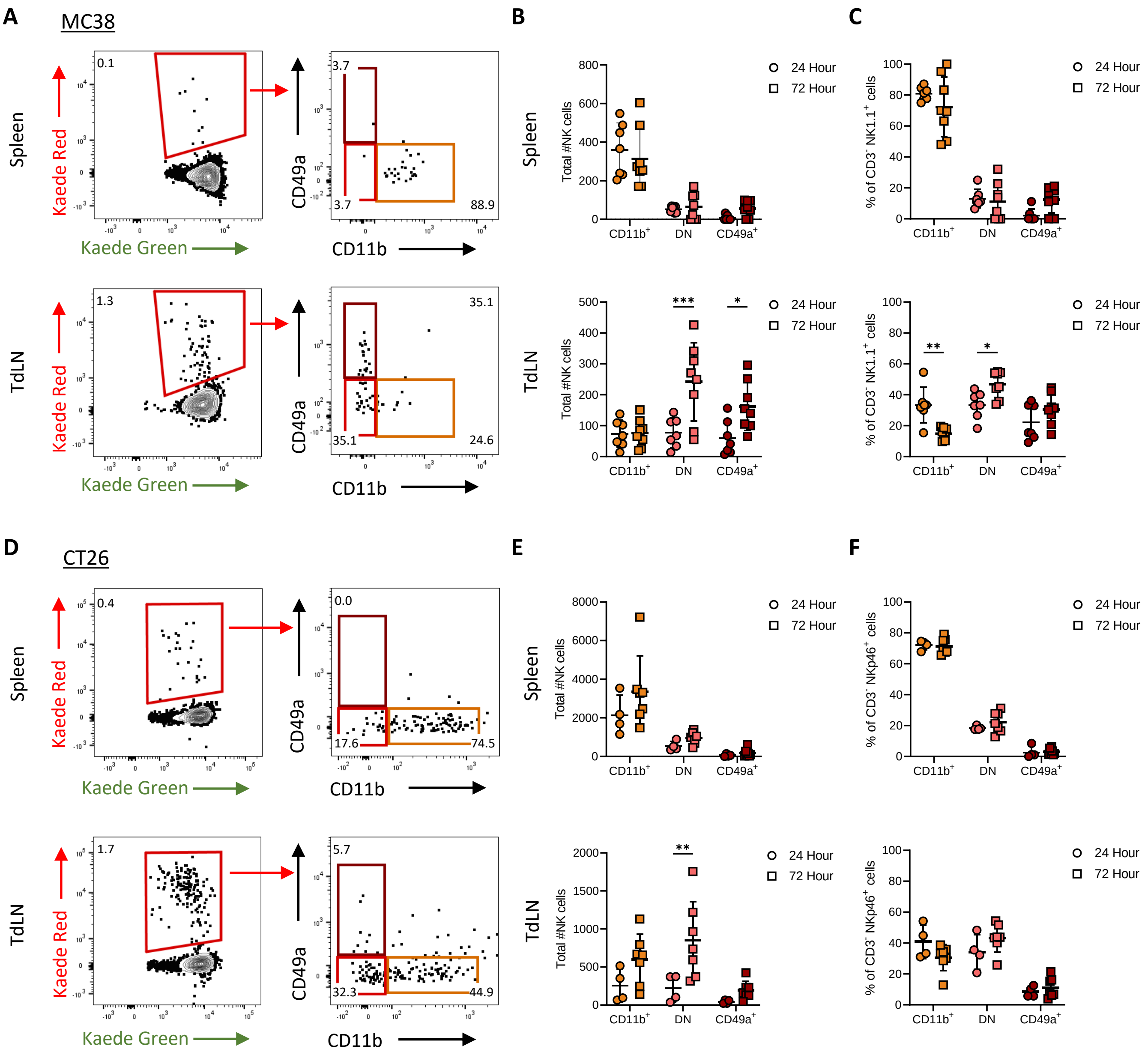
A
MC38
Spleen
TdLN
Kaede Red
Kaede Green
CD49a
CD11b
0.1
3.7
3.7
88.9
1.3
35.1
35.1
24.6
B
C
Spleen
TdLN
Total #NK cells
% of CD3- NK1.1+ cells
CD11b+
DN
CD49a+
24 Hour
72 Hour
***
*
**
*
D
CT26
0.4
0.0
17.6
74.5
1.7
5.7
32.3
44.9
E
F
% of CD3- NKp46+ cells
**

#### Fig S6.

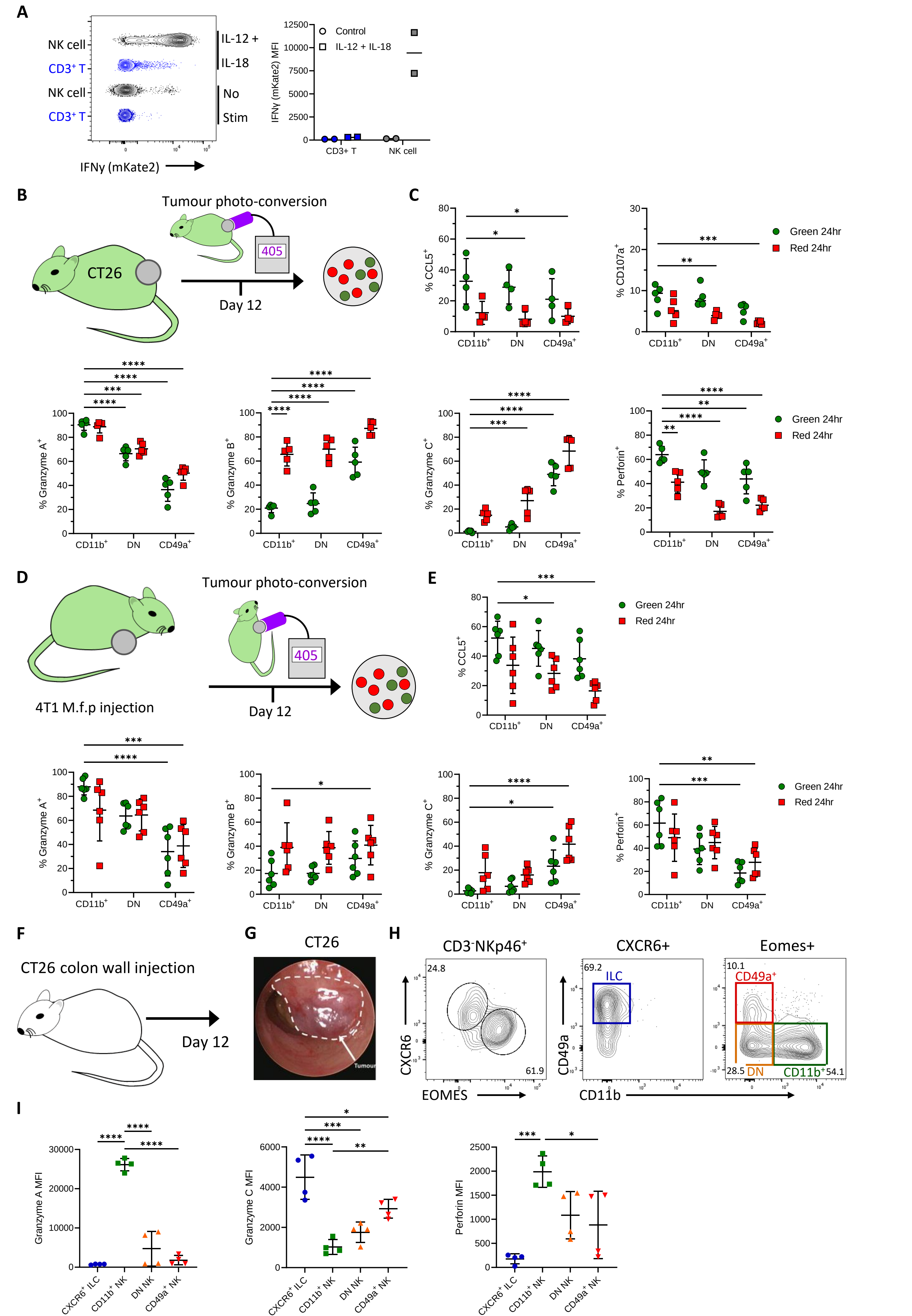

A
NK cell
CD3+ T
IL-12 +
IL-18
No
Stim
IFNγ (mKate2)
IFNγ (mKate2) MFI
Control
IL-12 + IL-18
CD3+ T
NK cell
B
Tumour photo-conversion
CT26
405
Day 12
C
Green 24hr
Red 24hr
% CCL5+
% CD107a+
% Granzyme A+
% Granzyme B+
% Granzyme C+
% Perforin+
CD11b+
DN
CD49a+
D
4T1 M.f.p injection
E
F
CT26 colon wall injection
G
CT26
Tumour
H
CD3-NKp46+
CXCR6+
Eomes+
CXCR6
EOMES
CD49a
CD11b
ILC
24.8
61.9
69.2
10.1
28.5
54.1
DN
CD11b+
I
Granzyme A MFI
Granzyme C MFI
Perforin MFI
CXCR6+ ILC
CD11b+ NK
DN NK
CD49a+ NK

Fig S7.

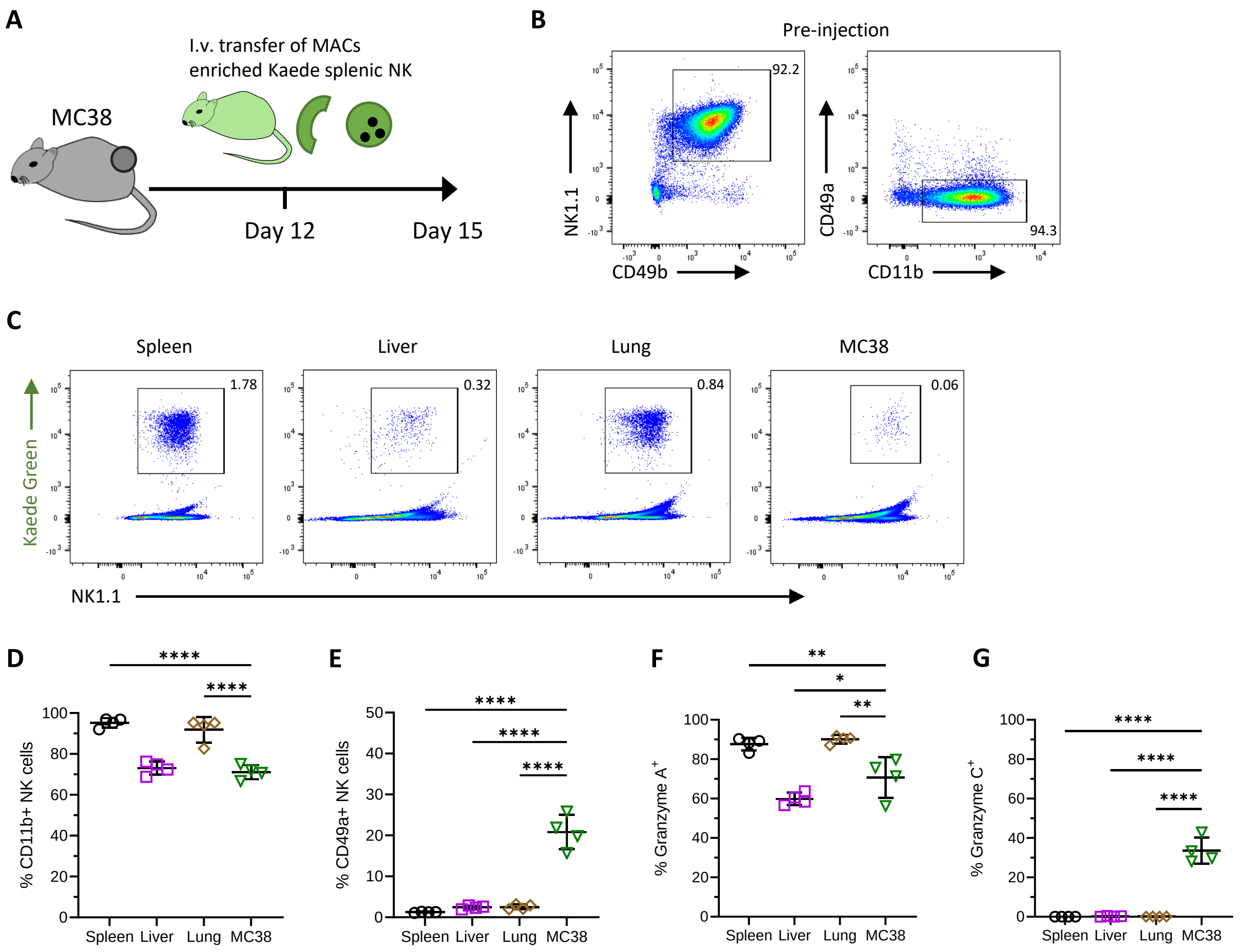
A
I.v. transfer of MACs
enriched Kaede splenic NK
MC38
Day 12
Day 15
B
Pre-injection
92.2
94.3
NK1.1
CD49b
CD49a
CD11b
C
Spleen
Liver
Lung
MC38
1.78
0.32
0.84
0.06
Kaede Green
NK1.1
D
% CD11b+ NK cells
E
% CD49a+ NK cells
F
% Granzyme A+
G
% Granzyme C+
Spleen Liver Lung MC38
****
**
*

#### Fig S8.

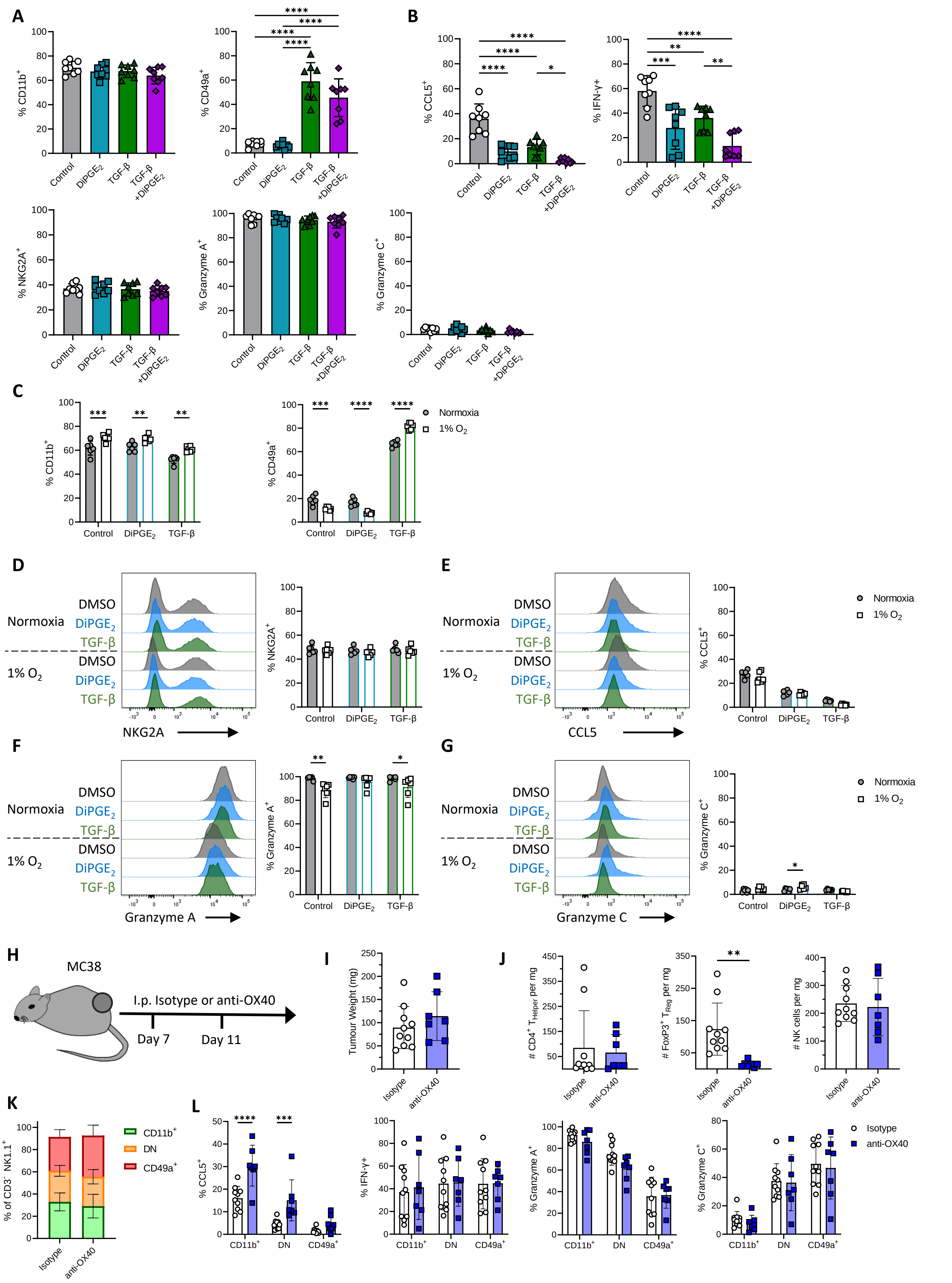

A
B
C
D
E
F
G
H
I
J
K
L
% CD11b+
% CD49a+
% CCL5+
% IFN-γ+
% NKG2A+
% Granzyme A+
% Granzyme C+
Control
DiPGE2
TGF-β
TGF-β +DiPGE2
Normoxia
1% O2
DMSO
NKG2A
CCL5
Granzyme A
Granzyme C
MC38
I.p. Isotype or anti-OX40
Day 7
Day 11
Tumour Weight (mg)
### CD4+ THelper per mg
### FoxP3+ TReg per mg
### NK cells per mg
Isotype
anti-OX40
% of CD3- NK1.1+
CD11b+
DN
CD49a+

**Fig S9.**

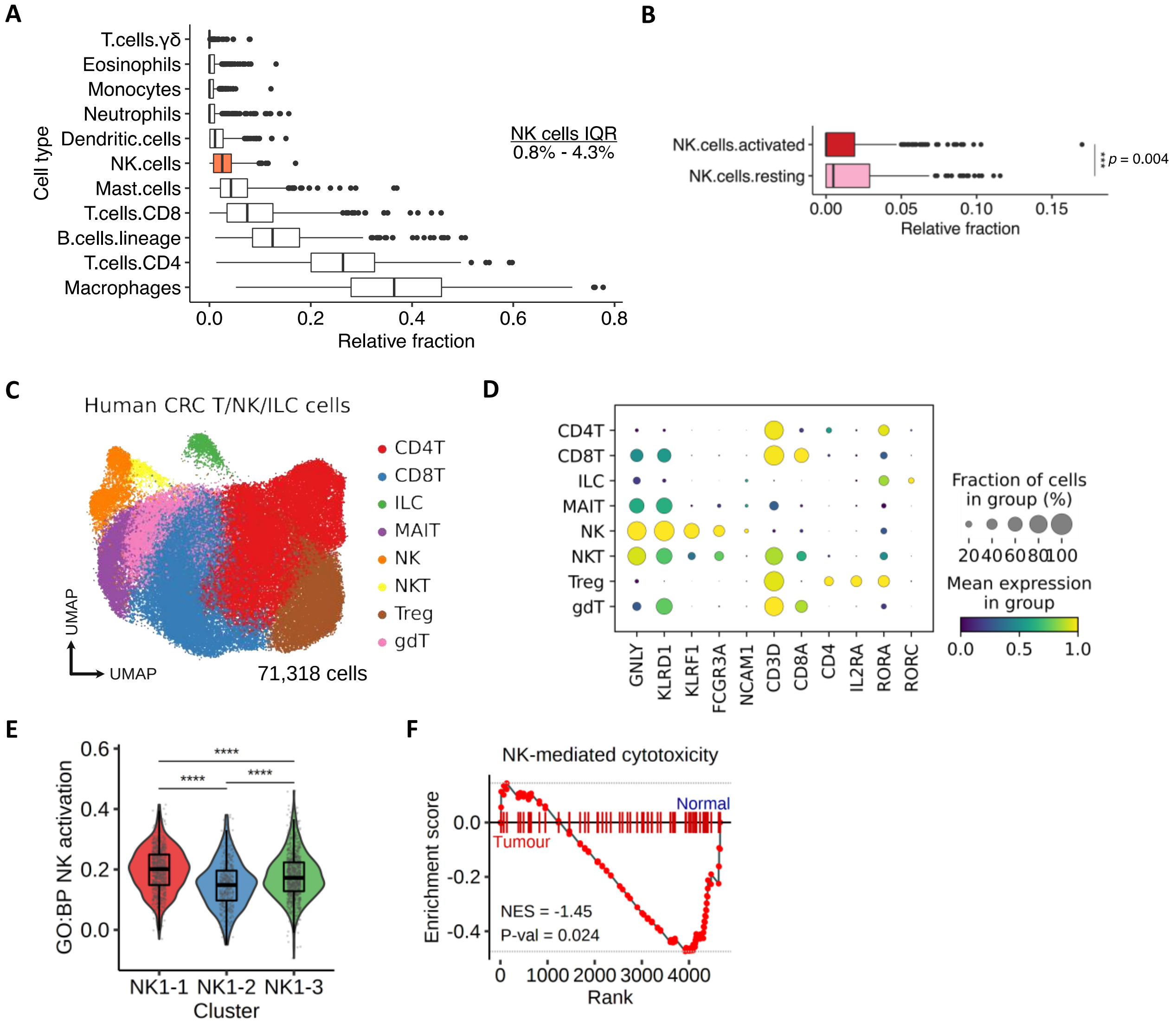
A
T.cells.γδ
Eosinophils
Monocytes
Neutrophils
Dendritic.cells
NK.cells
Mast.cells
T.cells.CD8
B.cells.lineage
T.cells.CD4
Macrophages
Cell type
0.0 0.2 0.4 0.6 0.8
Relative fraction
NK cells IQR
0.8% - 4.3%
B
NK.cells.activated
NK.cells.resting
p = 0.004
0.00 0.05 0.10 0.15
Relative fraction
C
Human CRC T/NK/ILC cells
CD4T
CD8T
ILC
MAIT
NK
NKT
Treg
gdT
UMAP
UMAP
71,318 cells
D
CD4T
CD8T
ILC
MAIT
NK
NKT
Treg
gdT
GNLY
KLRD1
KLRF1
FCGR3A
NCAM1
CD3D
CD8A
CD4
IL2RA
RORA
RORC
Fraction of cells in group (%)
20 40 60 80 100
Mean expression in group
0.0 0.5 1.0
E
GO:BP NK activation
0.0 0.2 0.4 0.6
****
****
****
NK1-1 NK1-2 NK1-3
Cluster
F
NK-mediated cytotoxicity
Enrichment score
0.0 -0.2 -0.4
Normal
Tumour
NES = -1.45
P-val = 0.024
0 1000 2000 3000 4000
Rank

**Fig S10.**

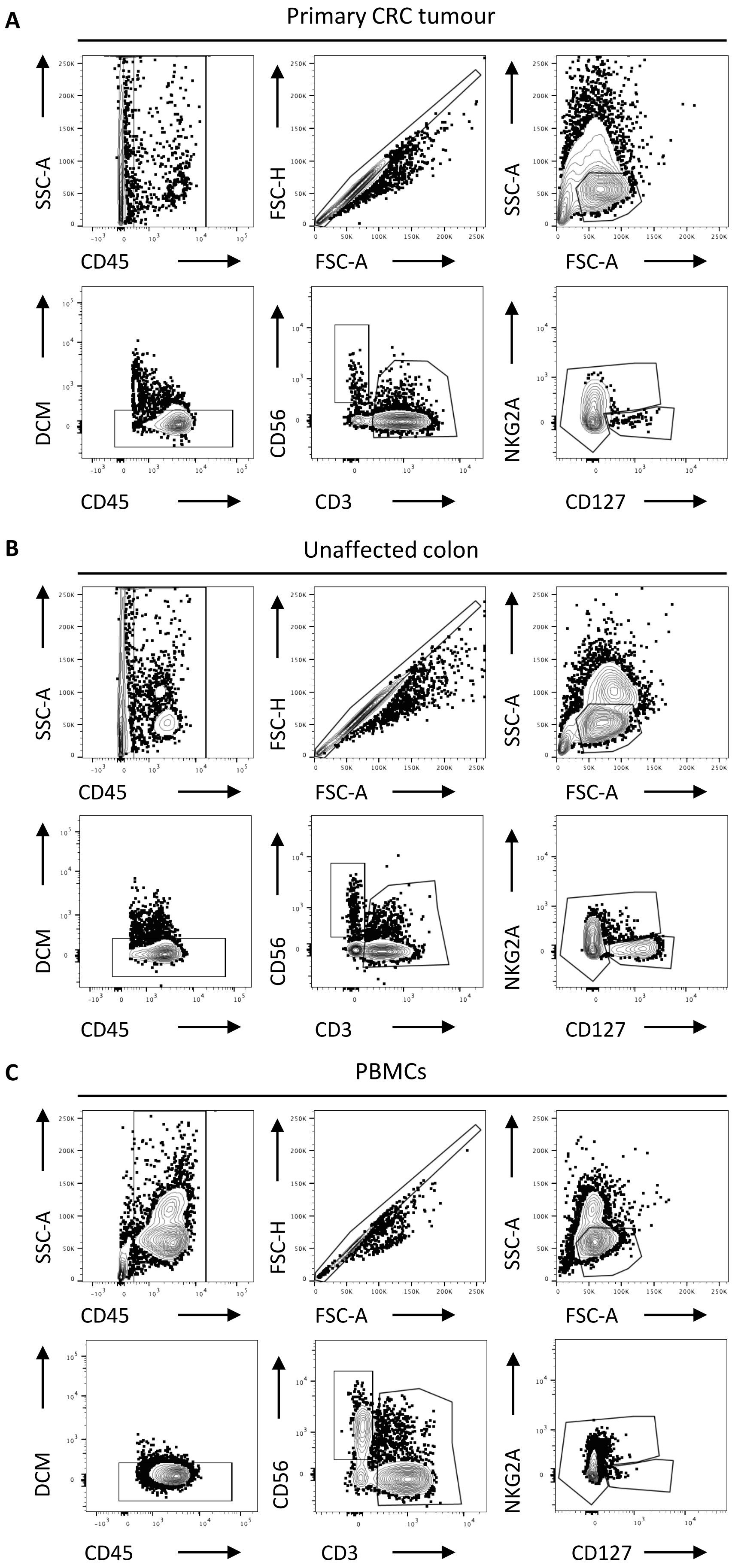
A
Primary CRC tumour
SSC-A
CD45
FSC-H
FSC-A
SSC-A
FSC-A
DCM
CD45
CD56
CD3
NKG2A
CD127
B
Unaffected colon
SSC-A
CD45
FSC-H
FSC-A
SSC-A
FSC-A
DCM
CD45
CD56
CD3
NKG2A
CD127
C
PBMCs
SSC-A
CD45
FSC-H
FSC-A
SSC-A
FSC-A
DCM
CD45
CD56
CD3
NKG2A
CD127

**Fig S11.**

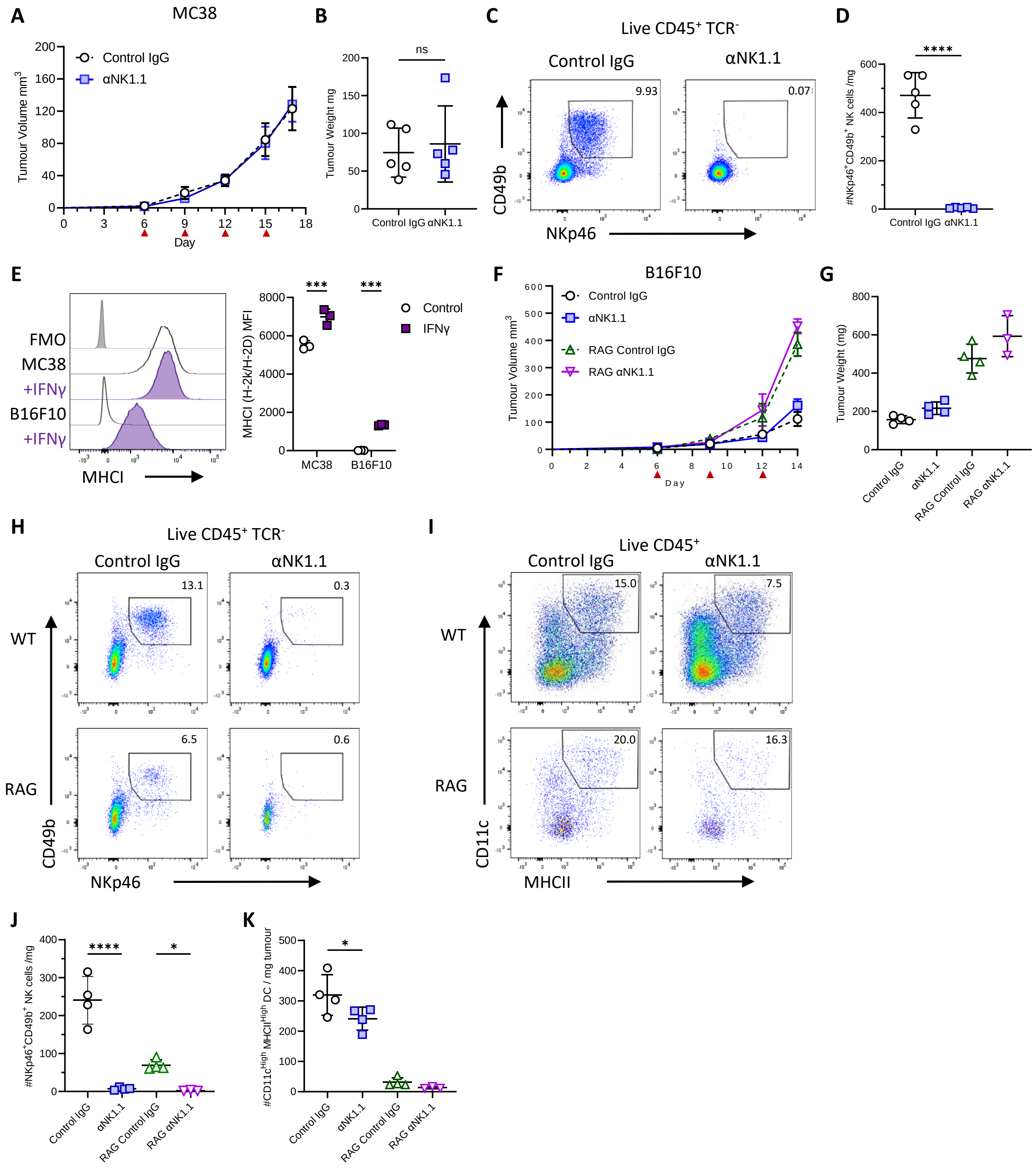

A
MC38
Control IgG
αNK1.1
Tumour Volume mm3
Day
B
ns
Tumour Weight mg
Control IgG
αNK1.1
C
Live CD45+ TCR-
Control IgG
αNK1.1
9.93
0.07
CD49b
NKp46
D
****
#NKp46+CD49b+ NK cells /mg
Control IgG
αNK1.1
E
FMO
MC38
+IFNγ
B16F10
+IFNγ
MHCI
***
***
Control
IFNγ
MHCI (H-2k/H-2D) MFI
MC38
B16F10
F
B16F10
Control IgG
αNK1.1
RAG Control IgG
RAG αNK1.1
Tumour Volume mm3
Day
G
Tumour Weight (mg)
Control IgG
αNK1.1
RAG Control IgG
RAG αNK1.1
H
Live CD45+ TCR-
Control IgG
αNK1.1
13.1
0.3
6.5
0.6
WT
RAG
CD49b
NKp46
I
Live CD45+
Control IgG
αNK1.1
15.0
7.5
20.0
16.3
WT
RAG
CD11c
MHCII
J
****
*
#NKp46+CD49b+ NK cells /mg
Control IgG
αNK1.1
RAG Control IgG
RAG αNK1.1
K
*
#CD11cHigh MHCIIHigh DC / mg tumour
Control IgG
αNK1.1
RAG Control IgG
RAG αNK1.1

**Fig S12.**

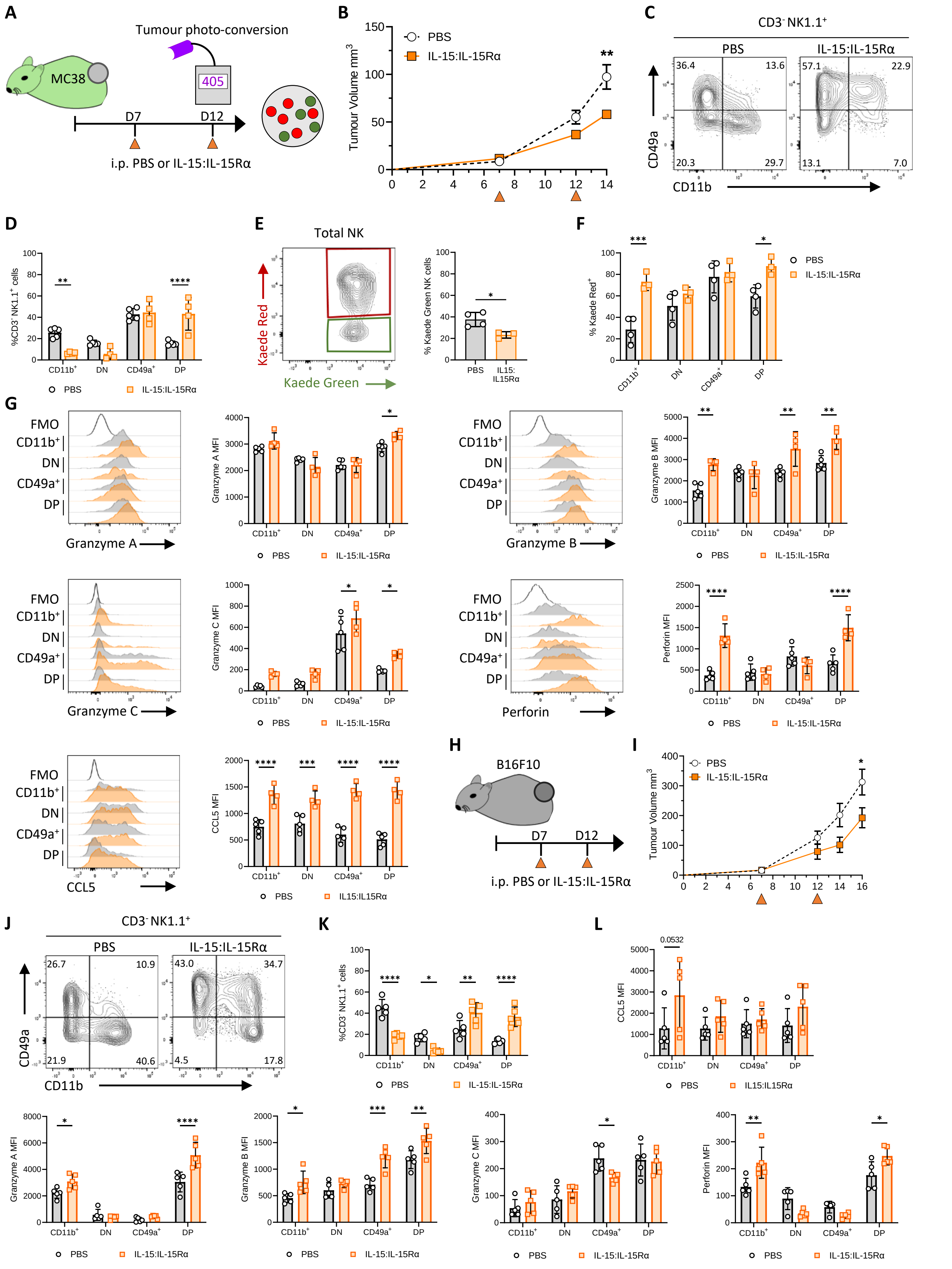
A
Tumour photo-conversion
405
MC38
D7
D12
i.p. PBS or IL-15:IL-15Rα
B
PBS
IL-15:IL-15Rα
Tumour Volume mm3
C
CD3- NK1.1+
PBS
IL-15:IL-15Rα
CD49a
CD11b
D
%CD3- NK1.1+ cells
CD11b+
DN
CD49a+
DP
E
Total NK
Kaede Red
Kaede Green
% Kaede Green NK cells
IL15:IL15Rα
F
% Kaede Red+
G
FMO
Granzyme A
Granzyme A MFI
Granzyme B
Granzyme B MFI
Granzyme C
Granzyme C MFI
Perforin
Perforin MFI
CCL5
CCL5 MFI
IL15:IL15Rα
H
B16F10
I
J
K
L
